# Antagonizing the serotonin receptor HTR2B drives antigen-specific cytotoxic T-cell responses and controls colorectal cancer growth

**DOI:** 10.1101/2025.02.26.640476

**Authors:** Surojit Karmakar, Girdhari Lal

## Abstract

Neuroimmune communication is known to control various regulatory systems in the body. Serotonin (also known as 5-hydroxytryptamine, HT) is one of the neurotransmitters produced predominantly by enterochromaffin cells in the gut and the neurons of the central nervous system. Immune and cancer cells express several HT receptors (HTRs), and how HTRs modulate various immune pathways and contribute to cancer growth and metastasis is unclear. RNA-seq data analysis shows that human colon adenocarcinoma tissues with high HTR_2B_ expression are associated with poor anti-tumor immune response and poor patient survival. Using an orthotopic mouse model of tumors, we show that HTR_2B_ and other serotonergic molecule expressions are positively associated with colon tumor progression. Antagonizing the HTR_2B_ using specific chemical antagonists significantly suppresses the progression of colon, breast, and melanoma tumors and metastasis in mice by promoting antigen-specific cytotoxic CD8 T cell response and downregulating the PD-L1 expression on tumor cells. Further, treatment with a combination of suboptimal doses of HTR_2B_ antagonists and suboptimal doses of immunotherapy (anti-PD1, anti-PD-L1, or anti-CTLA4 mAb) or chemotherapeutic drugs (5-fluorouracil, oxaliplatin, or irinotecan) showed a robust anti-tumor response and inhibited colon tumor growth. Our finding showed that antagonizing the HTR_2B_ holds a potent therapeutic advantage as a combinatorial regimen of either immunotherapy or chemotherapy to treat colon cancer.

**One sentence summary:** HTR_2B_ expression is associated with colon tumor progression, and antagonizing the HTR_2B_ inhibits tumor growth and metastasis.

## Introduction

Neurotransmitters and associated signals regulate the immune response through neuro-immune pathways (1). Serotonin (5-Hydroxytryptamine; HT) is an amine neurotransmitter regulating the immune response to various diseases, including cancer (*2*). Serotonin is primarily synthesized by the enterochromaffin cells of the gut from the essential amino acid L-tryptophan by the tryptophan hydroxylase I (TPH1) enzyme (*3*). Platelets and several other immune cells take up serotonin from the gut and distribute them throughout the body (*4*). Serotonin signals through fifteen different receptors, most belonging to the G protein-coupled receptors (GPCRs) except HTR_3_, which forms a ligand-gated non-specific cation channel (*5*). In addition to its role as a neurotransmitter in the CNS and periphery, HT also works as a vasoconstrictor (*6*), regulates angiogenesis (*7*), controls bone density (*8*) and gastrointestinal motility (*9*), and regulates diverse metabolic functions like maintenance of glucose homeostasis and obesity (*10*). HTR_2B_ is particularly interesting among different receptor subtypes because of its widespread distribution in the CNS and periphery. HTR_2B_ is coupled with stimulatory G-protein G_q_/11 in the cytosolic phase of the receptor and initiates a cascade of signaling pathways within the cell (*10*).

HTR_2B_ is expressed widely on several immune cells, such as dendritic cells, macrophages, and T cells (*11, 12*). Serotonin signaling through HTR_2B_ modulates macrophage polarization towards the anti-inflammatory M2 phenotype and suppresses polarization towards the pro-inflammatory M1 phenotype (*13*). Szabo *et al.* showed that HTR_2B_ signaling promotes the maturation of monocytic DCs via the upregulation of CD80, CD86, and CD83 (*14*). However, serotonin signaling through HTR_2B_ also suppresses DC-mediated IFN-γ^+^Th1 and IL-17^+^ Th17 differentiation and promotes IL-4^+^ Th2 differentiation *ex vivo* (*15*). The direct effect of HTR_2B_ signaling on T cells, B cells, or NK cells has not been explored. On the other hand, HTR_2B_ signaling widely has a pro-proliferative, pro-metastatic role on cancer cells *in vivo* and *in vitro* (*16–20*). Overall, the effect of HTR_2B_ signaling on both immune cells and cancer cells indicates that this signaling may promote cancer by enhancing cancer cell proliferation or modulating immune cell functions to some extent, potentially supporting tumor progression. However, the exact role of HTR_2B_ signaling in the immune regulation of cancer is not well defined.

In this study, we investigate the role of serotonin in modulating the immune response to cancer and how we can exploit the serotonergic system to promote an anti-tumor immune response. We showed increased intratumoral HTR_2B_ expression was associated with poor effector anti-tumor immune response and overall survival in colorectal cancer patients. HTR_2B_ antagonist potentiates the antigen-specific cytotoxic CD8 T cell response and inhibits mouse colon tumor growth and metastasis. Furthermore, a combination of a suboptimal dose of HTR_2B_ inhibitor along with suboptimal doses of immuno and chemotherapy showed a strong inhibition of colon tumor growth.

## Results

### HTR_2B-_high expression negatively correlated with survival of colorectal adenocarcinoma (COAD) patients

To assess the association of the serotonergic system with cancer progression, the cancer genome atlas (TCGA) gene expression data were analyzed for the expression of various serotonin receptors in COAD patient’s colon tissues. Among all the HTRs expressed in COAD tumors, only HTR_2B_ was found to affect the survival of COAD patients significantly (**Fig 1A** and Fig S1). We have found that high intratumoral HTR_2B_ expression was significantly associated with poor survival in the COAD patient cohort (**Fig 1A**). Immunofluorescence analysis of COAD tumor tissue microarrays showed increased expression of HTR_2B_ and HT in the lamina propria (**Fig 1B**) and tumor (**Fig 1C**) compared to control samples. Increased HT expression was observed in the neuronal (neurofilament NF-A^+^ cells) and non-neuronal cells in the lamina propria and colon in COAD patients (**Fig 1C**), indicating both neuronal and non-neuronal sources of HT within the tumor. Lamina propria of COAD patients also showed an increased expression of TPH1, a key rate-limiting enzyme for HT biosynthesis, compared to the control tissues (Fig S1B), explaining the higher HT levels within the lamina propria. However, TPH1, monoamine oxidase A (MAO-A, enzyme that degrades HT), and HT transporter (SERT; SLC6A) gene expression in TCGA data showed no significant association with the survival of COAD patients (Fig S1C).

**Figure 1:**
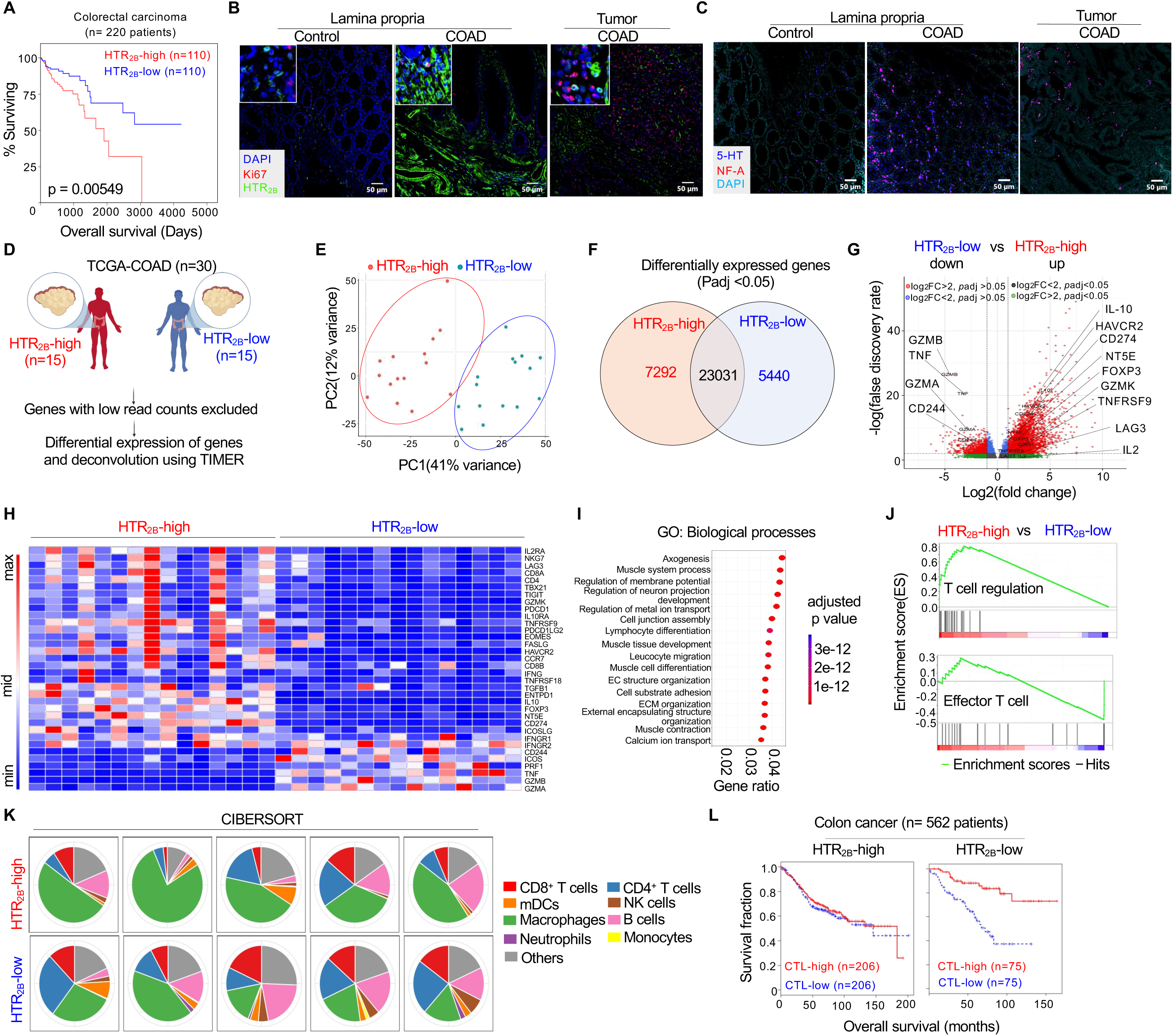
Dysregulated serotonergic system correlates with altered intra-tumoral immune composition in colon cancer patients. TCGA-COAD data were analyzed. **(A)** Kaplan-Meier survival analysis of HTR_2B_ expression (derived from TCGA data) in 220 colon adenocarcinomas patient’s cohort. “HTR_2B_-high” versus “HTR_2B_-low” data were segregated based on the cohort’s top or bottom 25% with intratumoral HTR_2B_ expression. Mantel-cox log rank test. **(B)** Representative images of HTR_2B_ and Ki67 in the control (lamina propria) and lamina propria of colon adenocarcinoma (COAD) tissue. The original magnification is 200X, and the inset images are 600X. **(C)** Representative images of serotonin (5-HT), neurofilament-A (NF-A) in the control (lamina propria) and lamina propria of colon adenocarcinoma (COAD) tissue IHC. Original magnification 200X. **(D)** RNA-seq datasets (n=30 samples) were segregated into “HTR_2B_-high” and “HTR_2B_-low” (highest fifteen and lowest fifteen intra-tumoral HTR_2B_ expression), and the differential expression of selected genes was analyzed. **(E)** Principal component analysis (PCA) of “HTR_2B_-high” versus “HTR_2B_-low” datasets (n=15 patients/group). **(F)** Venn diagram showing numbers of mutually exclusive or shared genes in each group. **(G)** A volcano plot shows differentially expressed genes (DEGs) of “HTR_2B_-high” versus “HTR_2B_-low” datasets (n=15 patients/groups) are shown. **(H)** Heatmap statistics differentially expressed selected gene sets. **(I)** Gene ontology of altered biological processes. **(J)** Gene set enrichment analysis (GSEA) of the curated gene sets that affect the regulatory and effector functions of the T cells. **(K)** Intra-tumoral immune cell signatures were analyzed by deconvolution using TIMER2.0 of HTR_2B_-high and -low datasets. **(L)** A TIDE computational analysis of the association between the tumor-infiltrating CD8 T cells (CTL) level (expression of CD8A, CD8B, GZMA, GZMB, and PRF1. Overall, patient survival with the intratumoral HTR_2B_ gene expression level. For each patient cohort, tumor samples were divided into HTR_2B_-high (samples with HTR_2B_ expression one standard deviation above the average, shown in the left survival plot) and HTR_2B_-low (remaining samples shown in the right survival plot) groups.

Further, to understand the effects of higher intra-tumoral HTR_2B_ on tumor progression, based on the highest (one standard deviation above the mode) or lowest (one standard deviation below the mode) HTR_2B_ expression, TCGA-COAD cancer patients were stratified, and the differential gene expression (DGE) analysis was performed (**Fig 1D**). The principal component analysis (PCA) of HTR_2B_-high and -low COAD patient cohorts formed two distinct clusters (**Fig 1E**), and DGE analysis (having adjusted *p-value* <0.05 and at least twofold change) showed that 7292 genes were upregulated in HTR_2B_-high cohorts, whereas 5440 genes upregulated in HTR_2B_-low cohorts (**Fig 1F**). Among all the differentially-expressed genes (DEGs), inflammatory genes such as GZMA, GZMB, CD244, and TNF-α were upregulated in the HTR_2B_-low cohort (**Fig 1G-1H**). In contrast, anti-inflammatory genes such as IL-10, HAVCR2, CD274, NT5E, FOXP3, TNFRSF9 and LAG3 were enriched in HTR_2B_-high cohort (**Fig 1G-1H**). Functional gene ontology (GO) analysis showed significant enrichment of various biological processes, cellular components, and molecular functions (**Fig 1I** and S1D-S1E). These GOs belong to leukocyte migration, lymphocyte differentiation, GPCR signaling in biological functions, integrin binding and cytokine signaling in molecular functions, and T-cell receptor complex in cellular components (**Fig 1I**, and S1D-S1E). Gene set enrichment analysis showed enriched genes associated with T cell effector function in the HTR_2B_-low cohort. In contrast, the HTR_2B_-high cohort had enriched gene sets related to T-cell suppression, angiogenesis, epithelial-mesenchymal transition, serotonin receptor signaling, serotonin transport, T-cell receptor signaling, cytokine-cytokine receptor interaction, chemokine receptor signaling and TGF-β receptor signaling (**Fig 1J**, and S1F). These data suggest that HTR_2B_-high and -low cohorts have differential immunological cellular and molecular signatures in the tumor microenvironment. Deconvolution of TCGA RNA-seq datasets using Tumor IMmune Estimation Resource (TIMER) 2.0 revealed that the HTR_2B_-high cohort had an enriched proportion of gene sets associated with macrophages and monocytes. In contrast, the HTR_2B_-low cohort had an enriched proportion of gene sets related to T, B, and NK cells (**Fig 1K**). Further, to identify any existing correlation between HTR_2B_ expression and cytotoxic CD8 T cell (CTL) response in COAD patients, the tumor immune dysfunction and exclusion (TIDE) platform was used. We found a significant dysfunctional CTL response in the HTR_2B_-high cohort compared to the HTR_2B_-low cohort (**Fig 1L**). These data suggest that an expression of lower anti-tumor CTL response genes is associated with HTR_2B_-high expression in COAD patients (**Fig 1L**). Together, we showed that higher intra-tumoral HTR_2B_ expression in COAD patients is associated with enriched immunosuppressive genes and a dysfunctional anti-tumor CTL response within the tumor microenvironment that might contribute to poor survival in HTR_2B_-high than HTR_2B_-low COAD patient cohorts.

### Antagonizing the HTR_2B_ signaling reduces the growth of colorectal and breast tumors in mice

To investigate further the significance of our findings related to HTR_2B_ in COAD patients’ tumor progression, we used a murine orthotropic colon cancer model. Mouse colon adenocarcinoma cell line MC38 cells were subcutaneously injected into the C57BL/6 mice **(Fig 2A**, left panel), and tumor growth was monitored **(Fig 2A**, middle panel). We found that tumor-bearing mice had significantly higher serum serotonin levels than naïve mice **(Fig 2A**, right panel). Tumor tissues showed expression of TPH1 and HT; interestingly, the expression of HTR_2B_ increased with tumor growth progression **(Fig 2B)**. To test the importance of HTR_2B_ signaling on tumor growth, C57BL/6 mice were subcutaneously injected with MC38 cells. From the 5^th^ day onward, mice were given daily intraperitoneal injections of either selective HTR_2B_-agonist (BW-723C86) or -antagonist (RS-127445), and tumor growth was monitored. Our data showed that HTR_2B_-agonist treatment significantly promoted tumor growth, whereas HTR_2B_-antagonist significantly reduced tumor growth in the immunocompetent mice (**Fig 2C**). Histopathology of tumor tissues from the HTR_2B_-agonist-treated mice displayed very few immune cell infiltrations, whereas HTR_2B_-antagonist-treated tumor tissues displayed enriched immune cell infiltration (**Fig 2D**). Since tumor growth was also associated with increased HT levels in the serum, we investigated the effects of circulating HT on tumor growth. To determine that, we used sertraline, a selective serotonin reuptake inhibitor (SSRI) that increases the free serotonin in the blood by blocking the serotonin transporter (SERT). Our data showed that sertraline treatment promoted the growth of colon tumors (**Fig S2A**). We also investigated the role of other HTRs on colon tumor growth by using specific antagonists. Our results showed that among the other HTR antagonists, such as ketanserin (pan-HTR_2_-antagonist) or SB-269970 (HTR_7_-antagonist) did not show any effect on tumor growth, whereas ritanserin (HTR_2A_-antagonist), or SB-215505 and RS-127445 (HTR_2B_-antagonist) significantly reduced tumor growth **(Fig 2E)**. However, HTR_2B_-antagonists had better anti-tumor activity than HTR_2A_-antagonists **(Fig 2E)**. To understand the broader anti-tumor effect of the HTR_2B_-antagonist, we also tested it on the tumor progression of mouse 4T1-luciferase breast cancer and B16F10 melanoma and metastasis of 4T1-luciferase breast cancer model. Our data showed that the HTR_2B_-antagonist inhibited melanoma **(Fig 2F)** and breast tumor growth **(Fig 2G)**. HTR_2B_-antagonist treatment also showed reduced breast cancer metastasis in the lung (**Fig 2H)**, as indicated by the reduced tumor-associated bioluminescence and in the lungs, with smaller and fewer tumor loci than the control **(Fig 2H)**. Together, we showed that HTR_2B_-signaling promotes tumor growth, and antagonizing this receptor has significantly reduced tumor growth and metastasis in different tissue-origin tumors.

**Figure 2:**
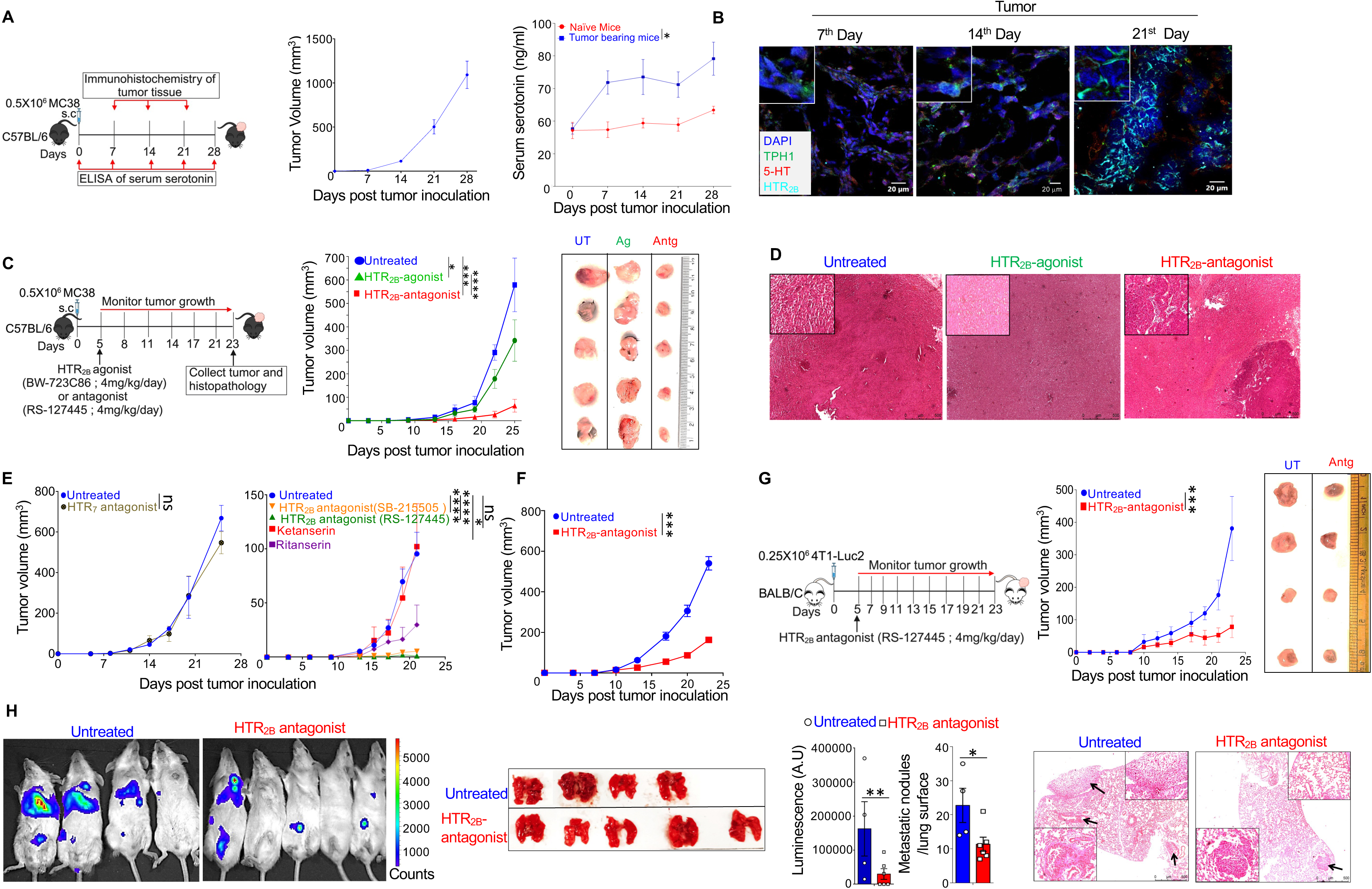
Antagonising the HTR_2B_ signaling reduces the growth of colorectal and breast tumors in mice. **(A**) C57BL/6 mice were subcutaneously injected with MC38 cells. The serum serotonin levels were measured at 0, +7, +14, and +21, +28-day post-tumor inoculation from the tumor-bearing and naïve mice using ELISA. **(B)** The **e**xpression of serotonin (red), TPH1 (green), and HTR_2B_ (cyan) was analyzed using immunofluorescence microscopy. The original magnification is 200X, and the inset images are 600X magnified. **C-D** C57BL/6 mice were subcutaneously injected with MC38 cells. 5^th^ day onward tumor injection, mice were treated with either HTR_2B_-agonist or HTR_2B_-antagonist or left untreated (n = 5 mice/group) for 21 days post-tumor inoculation. **(C)** Tumor growth (volume) kinetics and tumor images are shown. The data shown is representative of one of the four independent experiments. **(D)** Histopathology showing cellular infiltration within the tumor tissues. **(E)** C57BL/6 mice were subcutaneously injected with MC38 cells, and from the 5^th^ day onwards tumor injection, mice were treated with either HTR_7_ antagonist (SB-269970), HTR_2A_ antagonist (Ritanserin), pan-HTR_2_ antagonist (Ketanserin) HTR_2B_-antagonist [SB-215505 or RS-125505] or left untreated (n=6 mice/group). **(F)** C57BL/6 mice were subcutaneously injected with B16F10 cells. From the 5^th^ day of tumor injection, mice were treated with either HTR_2B_-antagonist or left untreated (n = 5 mice/group) for 25 days post-tumor inoculation. Tumor growth (volume) kinetics monitored. **(G)** Female BALB/c mice were injected with 4T1-Luc2 cells into the mammary fat pad and treated with either an HTR_2B_ antagonist or left untreated for 20 days (n = 5 mice/group). Tumor growth kinetics are shown. **(H)** Female BALB/c mice were intravenously injected into 4T1-Luc2 cells and were treated with either an HTR_2B_ antagonist or left untreated for 14 days (n=5 mice/group). Representative images showing quantification of the chemiluminescence (left panel). The images of the lungs are shown, and the number of tumor nodules of lung metastases is counted and plotted (middle panel), and lung histopathology images are shown (right panel). Two-way ANOVA with Sidak’s multiple comparison tests (A, C, E, F, G). Student’s ‘t’ test (H). p<0.05; *p<0.05; **p<0.01; ***p<0.001; ****p<0.0001. The error bar represents the ± standard error of means (SEM).

### A functional immune system is required for the anti-tumor activity of the HTR_2B-_ antagonist

To understand the mechanism of action of HTR_2B_ in tumors, we monitored the expression of HTR_2B_ on MC38 colon tumor cells and immune cells from the spleen and tumor. MC38 cells showed HTR_2B_ expression **(Fig 3A)**. Most immune cells, such as CD4, CD8, and γδ T cells in the tumor microenvironment of MC38 tumor-bearing mice, had higher HTR_2B_ expression than in the spleen **(Fig 3B)**. Since HTR_2B_ is expressed in both MC38 cells and immune cells, we further investigated the importance of HTR_2B_ in anti-tumor immunity. To test this, MC38 cells were subcutaneously injected into the NRG mice (NOD-RAG1^−/−^IL-2Rγ^−/−^; lack T, B, and NK cells) and treated with HTR_2B_-antagonist (RS-127445). Our data showed that HTR_2B_-antagonists failed to restrict tumor growth in the NRG mice **(Fig 3C)**, suggesting that functional effector immune cells are required for the anti-tumor function of HTR_2B_-antagonist. We further investigated whether exposure to HTR_2B_-antagonist in C57BL/6 mice is sufficient to generate activated immune cells capable of controlling tumor growth or whether tumor antigen is essential for HTR_2B_-antagonist-induced anti-tumor immunity. Thus, we subcutaneously injected C57BL/6 mice either with MC38 cells or sham injection. From the 5^th^ day onward, these mice were treated daily with HTR_2B_-antagonist for 10 days. On day 15 of the tumor injection, mice were euthanized, and lymph node cells were harvested from the tumor-draining lymph nodes. Lymph node cells were adoptively transferred into the NRG mice-bearing MC38 tumors. Interestingly, the lymph node cells exposed to only HTR_2B_-antagonists did not control the tumor growth in NRG mice **(Fig 3D)**. However, lymph node cells from the mice that received both MC38 cells and HTR_2B_-antagonist restricted tumor growth in the NRG mice **(Fig 3D)**. To understand whether HTR_2B_ antagonist treatment induced tumor-antigen-specific immune response, MC38 tumor-bearing mice were treated with HTR_2B_ antagonists and lymph node cells were isolated from the tumor-draining lymph nodes. These lymph node cells were then adoptively transferred into the NRG mice that received subcutaneous MC38 cells on one flank and B16F10 melanoma tumor cells on the contralateral flanks **(Fig 3E)**. Cells from the HTR_2B_-antagonist-treated MC38 tumor-bearing mice showed significant control of MC38 tumor growth but not the B16F10 tumor growth in the same NRG mice **(Fig 3E),** suggesting that the development of tumor antigen-specific immune response.

**Figure 3:**
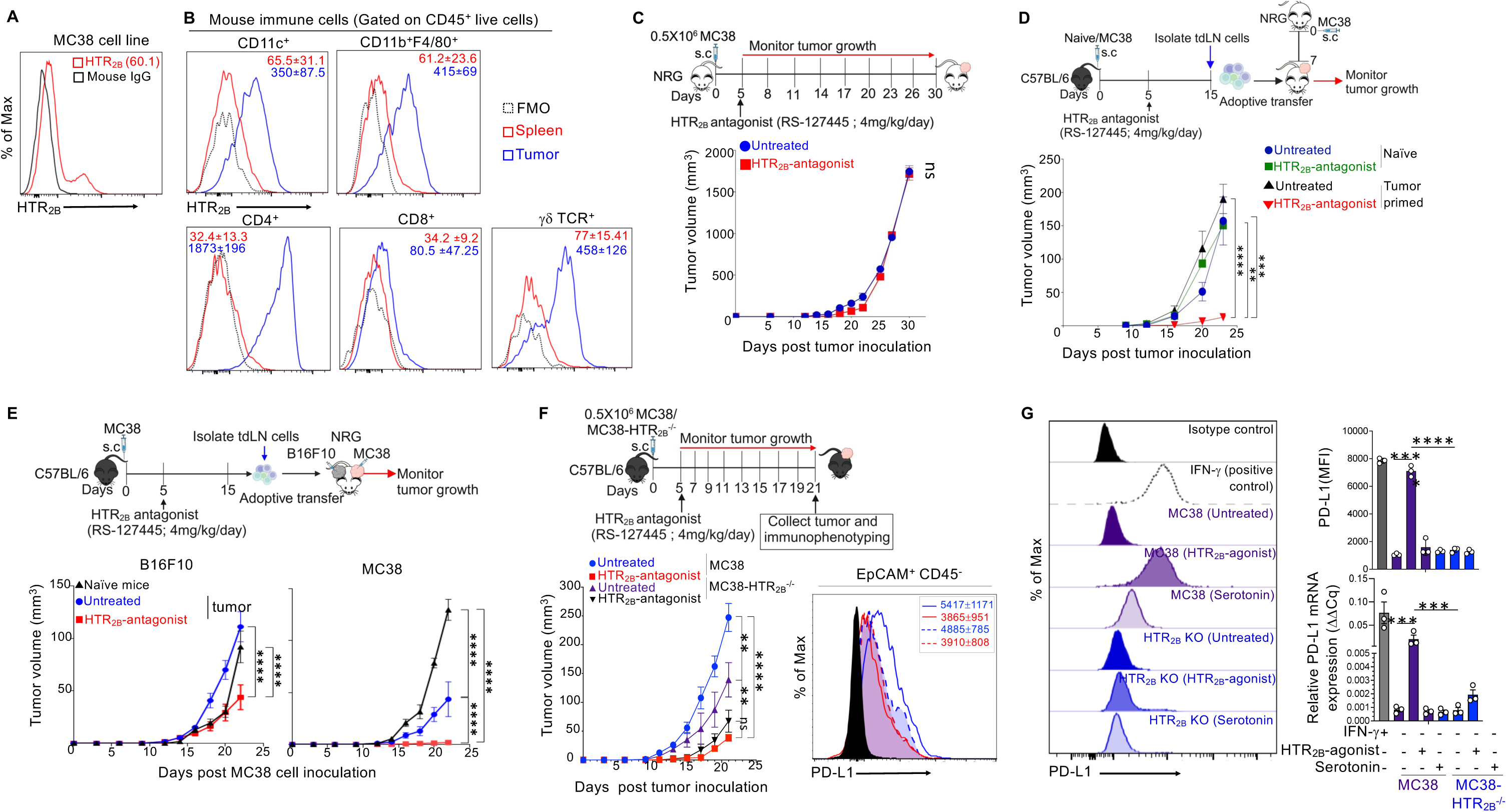
A functional immune system is required for the anti-tumor activity of the HTR_2B_-antagonist. **(A**) HTR_2B_ expression on the MC38 cell line was monitored using flow cytometry. Histogram overlay with the isotype control. (**B)** C57BL/6 mice were subcutaneously injected with MC38 cells, and after 21 days of tumor inoculation, HTR_2B_ expression in the different immune cell subsets from the spleen and tumor was monitored using flow cytometry. Representative histogram overlays are shown. The numbers mentioned in the histogram show the MFI of HTR_2B_ in the spleen and tumor. **(C)** RAG1^−/−^IL-2Rγ^−/−^ (NRG) mice were subcutaneously injected with MC38 cells, and 5^th^ day onward tumor injection, mice were treated with either HTR_2B_ antagonist or sham-treated (n = 5 mice/group) for 30 days post-tumor inoculation. Tumor growth (volume) kinetics are shown. N=2 independent experiments. **(D)** Total immune cells from the inguinal lymph nodes of naïve or MC38 tumor-bearing HTR_2B_ antagonist-treated or sham-treated C57BL/6 mice were adoptively transferred into the MC38 tumor-bearing NRG mice (n = 4 mice/group). Tumor growth kinetics are shown. **(E)** Total immune cells from the inguinal lymph nodes of HTR_2B_ antagonist treated or sham-treated MC38 tumor C57BL/6 mice were adoptively transferred into the NRG mice bearing B16F10 melanoma tumor in the left flank and MC38 tumor in the right flank (n = 5 mice/ group). Tumor growth kinetics of MC38 and B16F10 tumors were shown. **F-G** C57BL/6 mice were subcutaneously injected with MC38 or MC38-HTR_2B_^−/−^ cells, and 5^th^ day onward tumor inoculation, mice were treated with either HTR_2B_ antagonist or sham-treated (n=5 mice/group) for 21 days. On the 21^st^ day, the phenotype of tumor-infiltrating CD8 T cells was monitored. **(F)** Tumor volume kinetics are shown (bottom Left panel), and the histogram overlay shows PD-L1 expression on EpCAM^+^ CD45^−^ cancer cells (Bottom right panel). **(G)** MC38 or MC38-HTR_2B_^−/−^ cell lines were cultured in the presence or absence of HTR_2B_ agonist (BW-723C86; 50 μM) or serotonin (100 μM) for 72 hours. Histogram overlay and MFI of PD-L1 on the MC38 or MC38-HTR_2B_^−/−^ cells cultured in the presence or absence of HTR_2B_ agonist or serotonin (Right upper panel) and the relative expression of PD-L1 mRNA in MC38 or MC38-HTR_2B_^−/−^ cells following HTR_2B_ agonist or serotonin (Right lower panel). The values within the plot indicate MFI ± SEM. Two-way ANOVA with Sidak’s multiple comparison test (C, D, E, F). One-way ANOVA with Tukey’s multiple comparisons (H). *p<0.05; *p<0.05; **p<0.01; ***p<0.001; ****p<0.0001. The error bar represents the ± SEM.

To delineate the effects of intrinsic HTR_2B_ signaling in the cancer cells on the modulation of anti-tumor CD8 T cell response, MC38-HTR_2B_^−/−^ cells were generated by transfecting MC38 cell lines with SR-2B KO CRISPR/Cas9 plasmid **(Fig S2B)**. Then, we subcutaneously injected C57BL/6 mice with HTR_2B_ deficient or sufficient MC38 cells and treated them with HTR2B antagonists. We observed that the HTR_2B_^−/−^ on the MC38 tumor showed slower growth than the HTR_2B_^+/+^ tumor. **(Fig 3F)**. It has been reported that colon tumor cells in the microenvironment have higher expression of immune checkpoints like PD-L1 that suppress tumor-specific T cell response (*21–24*), an excellent target for immunotherapy in colon tumors (*22, 25, 26*). To test whether the deficiency of HTR_2B_ affected PD-L1 expression in the tumor microenvironment, single-cell suspensions from the tumor were stained for markers (EpCAM^+^CD45^−^ cells) to identify intratumoral MC38 cells and analyzed for PD-L1 expression using flow cytometry. Our data showed that a deficiency of HTR_2B_ reduced the PD-L1 on MC38 cells in the tumor microenvironment **(Fig 3G)**. To further validate whether the alteration of PD-L1 expression on MC38 cells is specific to HTR_2B_ signaling, HTR_2B_^−/−^ and HTR_2B_^+/+^ cells were *in vitro* stimulated with HT or HTR_2B_-agonist and expression of PD-L1 mRNA and protein were monitored. Our data showed that specific agonism of HTR_2B_ promotes PD-L1 expression in MC38 cells **(Fig 3H)**, similar to activation with cytokine IFN-γ(*27*). Together, these results suggest that the source of tumor antigen is required to generate an HTR2B-antagonist-driven functional anti-tumor immune response to control tumor growth.

### The HTR_2B_-antagonist-driven anti-tumor response is dependent on the CD8 T cells

Our data from patient cohorts showed that HTR_2B_-low COAD tumors had increased genes associated with cytolytic and effector function of CD8 T cells compared to HTR_2B_-high tumors (**Fig 1G, 1H,** and **1K**). Interestingly, HTR_2B_-high COAD tumors had dysfunctional cytotoxic CD8 T cell response compared to HTR_2B_-low tumors (**Fig 1L**). Immunofluorescence staining of MC38 tumor tissues showed that HTR_2B_-agonist (BW723C86) treated mice had significantly lower CD8 T cell infiltration within the tumor compared to HTR_2B_-antagonist (RS-1275445) treated mice (**Fig 4A**). Serum cytokine analysis showed significantly increased levels of IFN-γ and TNF-α, and significantly reduced anti-inflammatory cytokine IL-10 compared to the control group **(Fig 4B)**. Further, HTR_2B_-antagonist treated mice tumor had significantly increased activated CD8 T cells (CD69^+^CD8^+^), and central (CD8^+^CD44^+^CCR7^+^) and effector (CD8^+^CD44^+^CCR7^−^) memory cells in the tumor compared to the control group (**Fig 4C**). CD8 T cells in the HTR_2B_-antagonist treated mice tumor also had significantly increased effector CD8 T cells (IFN-γ^+^Granzyme B^+/−^cells) and cytolytic CD8 T cells (IFN-γ^+^Grnzyme B^+^cells) compared to the control tumors (**Fig 4C**). To understand the importance of CD8 T cells in HTR_2B_-antagonist driven anti-tumor immune response, CD8 T cells were depleted in C57BL/6 mice using an anti-CD8α monoclonal antibody (Clone YTS 169.4; −3, +1, +5, +9, +13, +17 days with respect to MC38 cell injection), and these mice were injected with MC38 cells and treated with HTR_2B_ antagonist. Compared to the isotype control antibody, the CD8 T cell depletion with anti-CD8α antibody was validated in the various lymph nodes (**Fig S2C**). Depletion of CD8 T cells in the mice prevented the HTR_2B_-antagonist-driven inhibition of tumor growth (**Fig 4D**). To further evaluate the potency of HTR_2B_-antagonist-induced functional anti-tumor CD8 T cells, C57BL/6 mice were injected with MC38 cells and treated with HTR_2B_-antagonist. On the 15^th^ day of tumor induction, CD8 T cells were isolated from the tumor of these mice and adoptively transferred into the NRG mice having MC38 tumors. CD8 T cells isolated from the HTR_2B_-antagonist-treated mice inhibited tumor growth in NRG mice compared to the CD8 T cells isolated from tumor-bearing control mice (**Fig 4E**), indicating the generation of potent long-lived tumor-specific CD8 T cells. These data suggest that the HTR_2B_-antagonist induces potent and long-lived effector and cytolytic CD8 T cells responsible for their anti-tumor properties.

**Figure 4:**
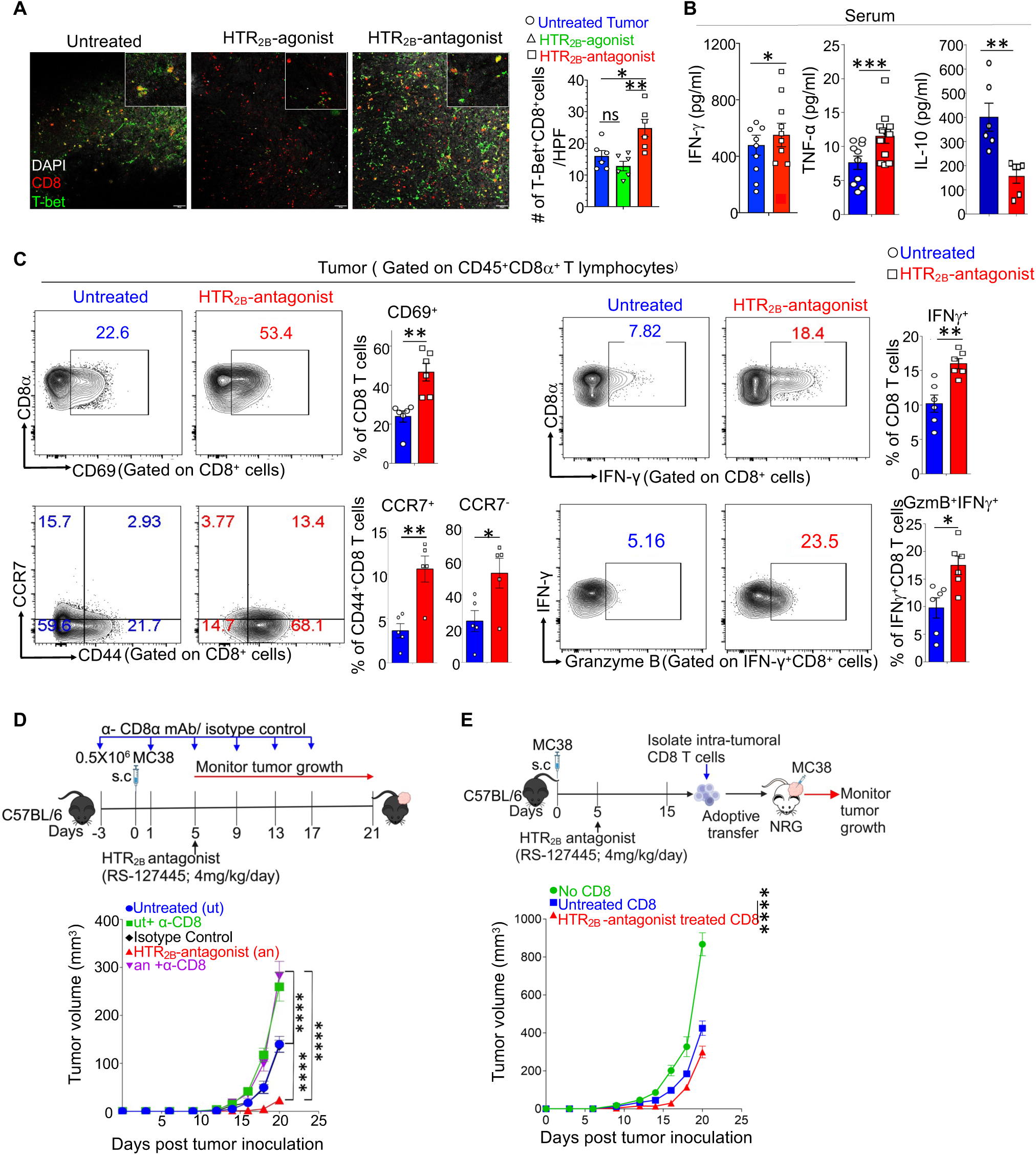
CD8 T cells play an essential role in HTR_2B_ antagonist-mediated reduction of tumor growth. **A-C.** C57BL/6 mice were subcutaneously injected with MC38 cells. 5th-day onward tumor injection, mice were treated with either HTR_2B-_agonist, HTR_2B-_antagonist, or sham-treated (n = 5 mice/group) for 21 days post-tumor inoculation. On the 21^st^ day, mice were euthanized, and immunophenotyping was performed from a single-cell suspension of tumor tissues. The serum cytokine levels were measured using ELISA. **(A)** Representative images of CD8 and T-bet in the untreated, HTR_2B_ agonist or antagonist-treated mice MC38 tumor tissue by immunohistochemistry (IHC). The original magnification is 200X, and the inset images are 600X. The right panel shows quantification and statistical comparisons of T-bet^+^CD8 T cells/ high power field (HPF) from the IHC images. **(B)** Serum TNF-α, IL-10, and IFN-γ levels were measured using ELISA from sham-treated or HTR_2B_ antagonist-treated MC38 tumor-bearing C57BL/6 mice. **(C)** Flow cytometric analysis of intra-tumoral CD8 T cell subsets. **(D)** C57BL/6 were injected with anti-CD8α monoclonal or isotype control antibody to deplete CD8 T cells. Mice were subcutaneously injected with MC38 cells and were treated with either HTR_2B_ antagonist or sham-treated (n=5 mice/ group). Tumor growth kinetics monitored. **(E)** Total intratumoral CD8 T cells from the naïve or MC38 tumor-bearing HTR_2B_ antagonist-treated or sham-treated C57BL/6 mice were adoptively transferred into the MC38 tumor-bearing NRG mice (n = 4 mice/group). One-way ANOVA with Tukey’s multiple comparison tests (A) Student’s ‘t’ test (B-C) Two-way ANOVA with Sidak’s multiple comparison test (D-E). *p<0.05; *p<0.05; **p<0.01; ***p<0.001; ****p<0.0001. The error bar represents the ± SEM.

### HTR_2B_-antagonist generates tumor antigen-specific CD8 T cells

CD8 T cells eliminate tumors by recognizing the specific tumor antigens through antigen-presenting cells (*28*). To investigate that HTR_2B_-antagonist drives the antigen-specific CD8 T cell response, the MC38 cell line was transfected with cytoplasmic ovalbumin expressing pCl-neo-cOVA plasmid (Addgene #25097)(*29*) to develop a cytoplasmic Ovalbumin-expressing MC38-cOVA cell line **(Fig S3A)**. This induced the expression of highly immunogenic chicken ovalbumin (OVA) protein in the MC38 cells that lacks a c-terminal secretory segment and forces the cells to present it through MHC class I instead of class II molecules and preferentially generate cOVA-specific CD8 T cells and, to some extent CD4 T cells via antigen cross-presentation (*29–31*). This allowed us to investigate the cOVA-specific CD8 T cell response using OVA-specific tetramers. C57BL/6 mice were subcutaneously injected with MC38-cOVA cells and treated with HTR_2B_ antagonist. On the 15^th^ day, the total intra-tumoral cells from both HTR_2B_-treated and untreated mice were harvested, labeled with CFSE dye, and restimulated *ex vivo* in the presence or absence of ovalbumin protein **(Fig 5A)**. Our results showed that CD8 T cells from the HTR_2B_-antagonist treated condition had a significantly higher proliferation rate after restimulation than control groups **(Fig 5A)**. Further, analysis of these CD8 cells using MHC class I (H2K^b^)-ova (SIINFEKL) tetramer (OT-I tetramer) showed that even though the HTR_2B_-antagonist did not enhance the frequencies of total cOVA-specific CD8 T cells within the tumor (**Fig 5B**), it enhanced the frequencies of CD69^+^ activated and IFN-γ^+^granzyme B^+^ effector cOVA-specific CTLs within the tumor **(Fig 5B)**. The overall IFN-γ and granzyme B expression were also higher in the tetramer^+^ CD8 T cells **(Fig 5C)**. These data indicate that HTR_2B_-antagonist treatment improved antigen-specific CTL response. To further elaborate on the phenotypes of these ova-specific CD8 T cells, OT-I mice having transgenic TCR on CD8 T cells to specifically recognize cOVA-peptide (SIINFEKL) (*32*) were subcutaneously injected with MC38-cOVA cells and treated with HTR_2B_ antagonist for 10 days. Then, the CD8 T cells from the tumor-draining lymph nodes (tdLN) were purified and tagged with Cell-trace violet (CTV) and adoptively transferred into the MC38-cOVA tumor-bearing PTPRC mice (CD45.1 congenic) **(Fig 5D)**. After five days, the tumor and lymph nodes of the PTPRC mice were harvested, and the proliferation and phenotypes of the transferred OT-I CD8^+^ T cells were monitored using flow cytometry. Our results showed that HTR_2B_ antagonist-treated OT-I CD8 T cells showed significantly higher proliferation within the congenic host compared to CD8 T cells from untreated mice **(Fig 5D)**. The immunophenotyping of the cells transferred from the antagonist-treated mice showed significantly higher frequencies of IFN-γ^+^granzyme B^+^ cells and lower frequencies of CTLA4^+^ or PD1^+^ or terminally exhausted (PD1^+^Tim3^+^Lag3^+^) cells **(Fig 5E)**, indicating an improved cytotoxic and reduced regulatory capacity in those tumor-specific CD8 T cells. These observations show that HTR_2B_ antagonism generates potent and robust antigen-specific CD8 T cell responses in colon tumors.

**Figure 5:**
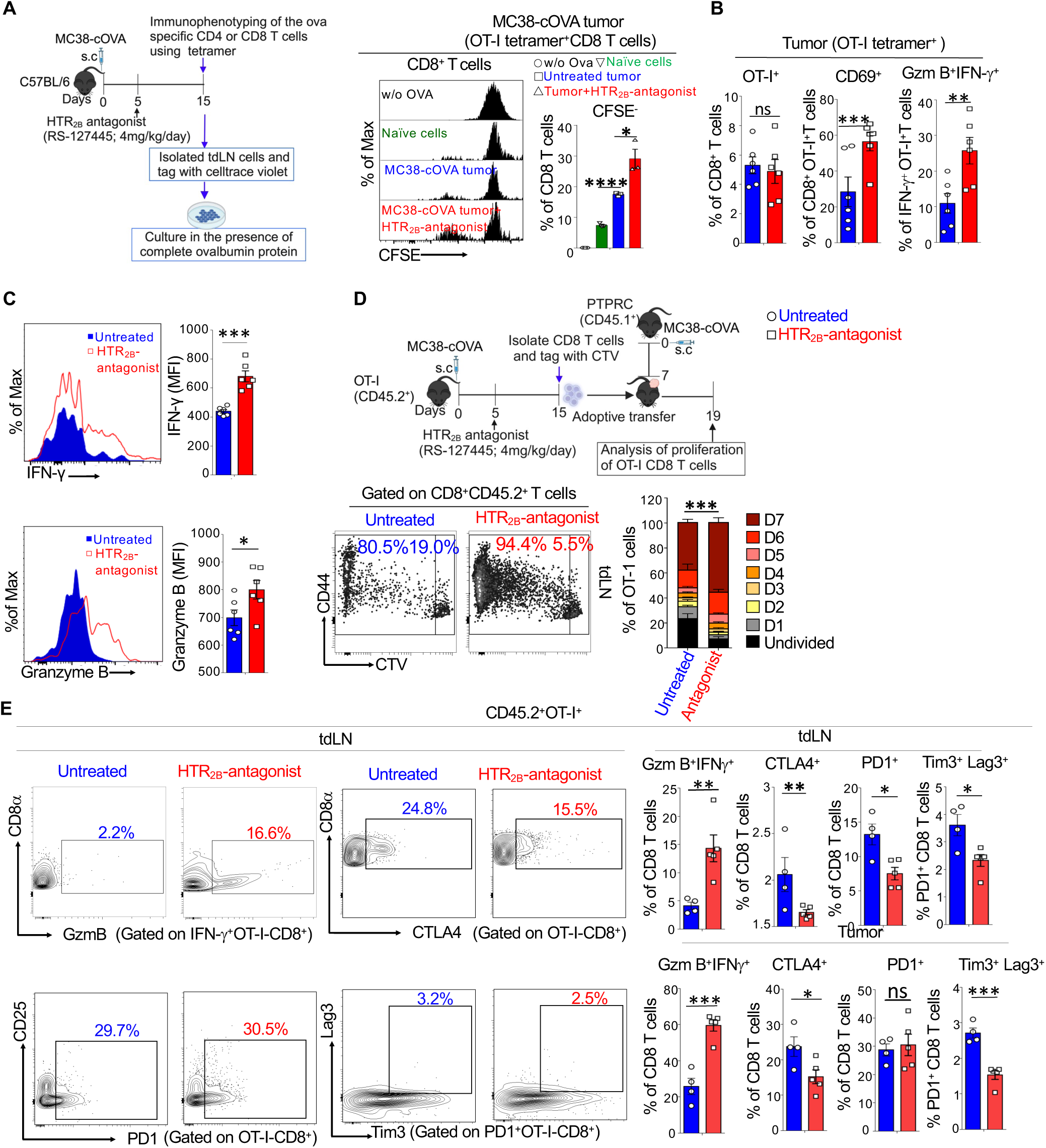
HTR_2B_ antagonist improves the antigen-specific CTL response in mice colorectal cancer model. **A-C.** C57BL/6 mice were injected with MC38-cOVA. They were either treated with HTR_2B_ antagonist or sham-treated (n = 5 mice/group). On the 15^th^ day, mice were euthanized, and total lymph node cells were tagged with CFSE and restimulated ex vivo with complete ovalbumin protein (50 μg/ml) for 5 days, or immunophenotyping was performed from the cells of tdLN and tumor. A stacked histogram and statistical analysis show the division of CFSE-tagged CD8 T restimulated *ex vivo.* **(B)** Flow cytometric analysis shows intra-tumoral OT-I tetramer^+^ total and different subsets of CD8 T cells. **(C)** Histogram overlay showing the expression of IFN-γ and granzyme B in intra-tumoral OT-I tetramer^+^ CD8 T cells from sham-treated or HTR_2B_ antagonist-treated mice. **D-E.** Cell-trace violet (CTV)-tagged CD45.2^+^OT-I CD8 T cells (1X10^6^ cells/mice) from MC38-cOVA tumor-bearing sham-treated or HTR_2B_ antagonist-treated OT-I mice were adoptively transferred into the MC38-cOVA tumor-bearing CD45.1^+^congenic mice. After 4 days, the division and phenotypes of the adoptively transferred cells (CD45.2^+^) were monitored. Representative flow cytometric plot and statistical analysis showing the division of CTV-tagged adoptively transferred OT-I-CD8 T cells (lower panel). **(E)** Flow cytometric analysis and the statistical comparisons of flow cytometric analysis of various subsets of adoptively transferred OT-I CD8 T cells from tumor and tdLN of PTPRC. One-way ANOVA using Tukey’s multiple comparison test (A, D) and Student’s ‘t’ test (B, C, E). ns=not significant; *p<0.05; **p<0.01; ****p<0.001; ****p<0.0001. Error bars represent mean ± SEM.

### Low doses of HTR_2B_-antagonists enhance the efficacy of suboptimal immuno- or chemotherapy in controlling colon tumor growth

Although immunotherapies such as immune checkpoint blockers (ICBs) or chemotherapeutic agents showed remarkable success rates, dose-associated adverse events limit their efficiency in the clinics (*33–35*). However, many patients resist the therapy due to poor immune infiltrations and immune evasion mechanisms. Since HTR_2B_-antagonist generates potent and durable tumor-antigen specific cytotoxic T cell response, we wanted to understand if we can combine the low dose of HTR_2B_-antagonist with suboptimal doses of immune checkpoint blocker antibodies or chemotherapeutic drugs currently used to treat COAD patients to improve their anti-tumor function with less toxicity. To determine the prognostic value of HTR_2B_ in the treated melanoma patients cohort, we checked the outcome of anti-PD1 therapy in previously published melanoma patients cohort (*36*) based on the high or low intratumoral HTR_2B_ expression. We found that high pre-treatment HTR_2B_ expression was strongly associated with poor overall survival of patients **(Fig 6A)**. To further validate it in a pre-clinical mouse model, C57BL/6 mice were subcutaneously injected with MC38 cells and treated with combinations of low doses of HTR_2B_-antagonists with suboptimal doses of either anti-PD1 (Clone RMP1-14), anti-PD-L1 (Clone 10f.9G2), or anti-CTLA4 mAbs (Clone 9H10) **(Fig 6B)**. Suboptimal doses of immune checkpoint blocker antibodies (50 μg/mice every 4^th^ day) were used over the preferred doses (200 μg/mice/4^th^ day) to reduce the high-dose-associated toxicities. Such low doses of antagonists significantly synergize with either anti-PD1, anti-PD-L1, or anti-CTLA4 antibodies and reduce tumor growth **(Fig 6B)**. The combined therapy of HTR_2B_-antagonist and anti-PD1 also significantly enhanced IFNγ^+^granzyme B^+^ cytotoxic effector CD8 T cell population within the tumors **(Fig 6C),** showing potent T cell response. Similar to the mAbs, HTR_2B_ antagonist was also used in a combinatorial regimen with selected FDA-approved first and second-line chemotherapeutic drugs used in COAD such as oxaliplatin (*37*), 5-fluorouracil (5-FU) (*38*) and irinotecan (*39*) to target cancer cell proliferation and generate potent anti-tumor immune response simultaneously. Like the ICBs, we used suboptimal doses of the chemotherapeutic agents to avoid dose-related toxicities in our pre-clinical colon tumor models **(Fig 6D)**. Results showed that the combinatorial approach also significantly enhanced the efficacies of suboptimal doses 5-FU, oxaliplatin, and irinotecan in pre-clinical settings of the MC38 tumor model **(Fig 6D)**. Altogether, our data showed that low doses of HTR_2B_-antagonists in combination with suboptimal doses of ICB antibodies or chemotherapeutic drugs give a better therapeutic advantage in controlling colon tumor growth with lower potential toxicities.

**Figure 6:**
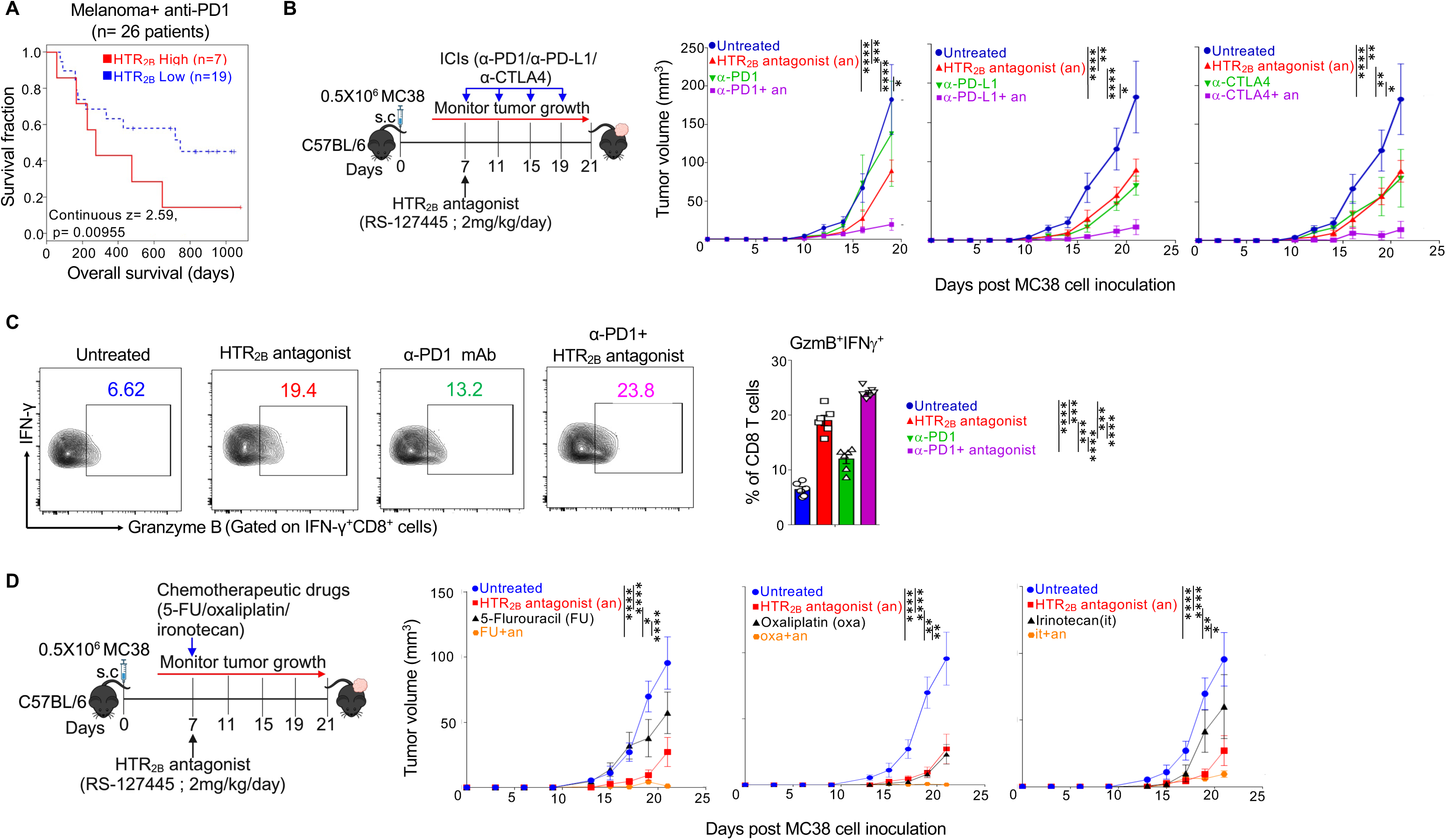
HTR_2B_ antagonist treatment enhances the efficacy of the immune checkpoint blockade antibodies and chemotherapeutic drugs on the MC38 tumor growth in mice. **(A)** The Kaplan Mayer survival analysis HTR_2B_ expression (derived from TCGA data) in 26 melanoma patients cohort treated with anti-PD1 ICB (nivolumab). z=2.59, P=0.00955. A positive Z score indicates that the expression of HTR_2B_ is negatively correlated with the therapeutic outcome, and the p-value indicates the significance level of the comparison between HTR_2B_-low and HTR_2B_ –high groups and calculated by two-sided Wald test in a Cox-PH regression model. **B-C.** C57BL/6 mice were injected subcutaneously with MC38 cells, and from 5^th^ day onward tumor injection, mice were treated with either immune checkpoint blockers (anti-PD1, anti-PD-L1, anti-CTLA4; 50 µg/mice every 4^th^ day) or HTR_2B_ antagonist (RS-127445; 2mg/kg/day) or a combination of both or sham-treated (n=5 mice/group) for 21 days post tumor inoculation. **(B)** Tumor volume kinetics of combination therapy using either anti-PD1, anti-PD-L1, or anti-CTLA4 mAb. **(C)** Flow cytometric and Statistical comparison of intra-tumoral effector (IFN-γ^+^ Granzyme B^+^) CD8 T cell population measured from untreated or treated with HTR_2B_ antagonist alone, anti-mouse anti-PD1 mAb alone, or a combination of both (anti-PD1 plus RS-127445). **(D)** C57BL/6 mice were injected subcutaneously with MC38 cells. From the 5^th^ day onward tumor injection, mice were treated with either chemotherapeutic drugs [5-Fluorouracil (4 mg/kg/day) or oxaliplatin (4 mg/kg/day for 5 days), or Irinotecan (5 mg/kg/day)] alone, or HTR_2B_ antagonist (RS-127445; 2 mg/kg/day) alone, or a combination of both or sham-treated (n=5 mice/group). **(H)** Tumor volume kinetics of combination therapy using either 5-FU or oxaliplatin alone or irinotecan. Two-way ANOVA, Sidak’s multiple comparison tests (B, D), and one-way ANOVA (C). *p<0.05; **p<0.01; ***p<0.00; ****p<0.0001. The error bar represents the mean ± SEM.

## Discussion

Different neurotransmitters affect the immune response to various diseases, including cancer, through neuro-immune communication. In the present study, we have identified a pathway through which serotonin influences tumor progression. Serotonin is a neurotransmitter that perturbs various functions in the brain and the peripheral organs in the body. Although many reports suggest the presence of various serotonin receptor subtypes in different immune cells subsets, such as macrophage, dendritic cells, and T cells, the detailed role of serotonin signaling in modulating their functions in homeostasis and diseases is not clearly understood. In the present study, we show that different components of the serotonergic system are present in various subsets of immune cells and cancer cells. Ling *et al.* reported that mice deficient in peripheral serotonin (TPH1^−/−^ mice) showed reduced MC38 tumor growth, which validates the role of peripheral serotonin in supporting colon tumor progression (*40*). In addition to colon tumors, HTR_2B_ antagonism has a similar impact on the primary tumor and the lung metastases formation in the 4T1 triple-negative breast cancer model, indicating conservation of the mechanism across various epithelial cancer types. HTR_2B_ antagonism reduces tumor progression by modulating the adaptive component of the anti-tumor immune response. Blocking the HTR_2B_ signaling exaggerates the proliferation and cytotoxic potential of the antigen-specific CD8 T cells within the tumor microenvironment. However, the detailed mechanism of HTR_2B_ signaling mediated alteration of antigen-specific cytotoxic CD8 T cell response remains to be further explored. In the present study, we could not use the HTR_2B_^−/−^ mice to validate the observed effects on the CD8 T cells due to their high mortality rates caused by defective heart valve development due to the deficiency of HTR_2B_ during developmental processes (*41, 42*).

After diagnosis of cancer, many patients suffer from mental health problems such as depression or anxiety and also show symptoms of major depressive episodes (*43–46*). Selective serotonin reuptake inhibitors (SSRIs) are one the most prescribed medications used for anxiety and depression (*47, 48*). Our results showed that serum serotonin levels, as well as levels of serotonin-synthesizing enzyme TPH1, increase with colon tumor growth, and mice treated with SSRI show enhanced tumors in mice. However, the incidence of death was prevalent with chronic SSRI treatment in breast, prostate, lung, and colon cancer patients (*49–51*). Recently, it has been shown that HTR_2B_ can be one of the diagnostic markers for various grades of colon cancer (*52*), and HTR_2B_, through CREB1-ZEB1 axis-mediated signaling, drives the metastatic(epithelial-mesenchymal transition) pathways (*53*). Interestingly, about 95% of serotonin is produced in the peripheral cells by enterochromaffin cells in the gut by the enzyme TPH1, whereas TPH2 drives the serotonin expression mostly in neurons (*11*). Mice deficient in TPH1 show less severe colitis and colon tumors (*54, 55*), and HT drives colon tumors’ tumor growth and angiogenesis. Targeted deletion of HTR_2B_, specifically in the enterocytes, enhanced gut inflammation and colon tumorigenesis (*56*). It is also reported that fluoxetine, an HT reuptake inhibitor treatment, promotes HT levels in the colon during inflammation-induced colitis development and ameliorates inflammation and further progression into colitis-associated cancer development (*57*), validating the anti-inflammatory potential of HT. HTR_2B_ gene deficiency or decreased expression was reported to have impulsivity in mice and humans (*58*), resistance to SSRI (*59*), sleep abnormalities, and hyperactivities in mice (*60*). Complete lack or deficiency of HTR_2B_ (HTR_2B_^−/−^) exacerbates weight loss with peripheral lipopolysaccharide injection, and mice have targeted deficiency of HTR_2B_ in microglial limits neuroinflammation in neonates and sickness behaviors in adulthood (*61*). Together, these studies point out that HTR_2B_ signaling has various immunoregulatory functions.

Many therapies, such as immune checkpoint inhibitors, target the anti-tumor cytotoxic CD8 T cell response to enhance tumor rejection. Despite their spectrum of success, their efficacy is still limited mainly due to inefficient immune activation and potent immune escape mechanisms by the tumor (*62–65*). In addition to the immune checkpoint inhibitors, different drugs are also used as chemotherapeutic agents to limit the division and proliferative potentials of the cancer cells within the tumor. These agents also sensitize cancer cells to immune cell-mediated killing. However, their success rate alone is also minimal due to the drug resistance mechanisms developed by the cancer cells(*66–68*). HTR_2B_ antagonist showed great potential in enhancing the CTL response against cancer, proving its potential as an anti-cancer therapeutic agent. In the preclinical *in vivo* mice model, combinations of lower doses of HTR_2B_ antagonists with the suboptimal doses of FDA-approved ICBs used in COAD, such as anti-PD1, anti-PD-L1, and anti-CTLA4 therapies or chemotherapeutic agents such as 5-FU, oxaliplatin and irinotecan in CRC significantly control the tumor growth. Due to the use of lower doses of the drugs, this approach not only reduces the chances of dose-related adverse events but also gives a tremendous advantage in utilizing ICB therapy with better efficacies and reduces the treatment cost. Our results strongly support the clinical development of therapeutic regimes targeting HTR_2B_ signaling in cancer and the development of combinatorial approaches using HTR_2B_ antagonists along with other therapeutic strategies that directly impact the cancer growth and the cytotoxic potentials of the CD8 T cells.

## Material and methods

### Cell line culture

Mouse colon adenocarcinoma cell line MC38 was a kind gift from Dr. Galina V. Shurin from the University of Pittsburgh. Mouse breast cancer cell lines 4T1-Luc2 and mouse melanoma cell line B16F10 were received from the National Centre for Cell Science, Pune. MC38 and B16F10 cells were cultured in complete Dulbecco’s modified Eagle media [DMEM supplemented with 10% fetal bovine serum, 2 mM glutamine, 0.1 mM MEM non-essential amino acid, 1 mM sodium pyruvate, 10 mM HEPES, 50 μg/ml antibiotics (gentamycin, penicillin, and streptomycin)]. The 4T1-Luc2 cell was cultured in complete RPMI-1640 media [RPMI supplemented with 10% FBS, 1 mM sodium pyruvate, 2 mM glutamine, and antibiotics]. All cell lines were maintained at 37°C in a 5% CO_2_ incubator. To treat the cell lines *in vitro,* all the adherent cells were plated in the tissue culture plates for 24 hours before the treatment with agonists or antagonists for indicated time points.

### 2.1 Mice experimental protocols

Six to eight weeks old C57BL/6J (RRID: IMSR_JAX:000664), BALB/c (RRID: IMSR_JAX:000651), NRG (NOD.Cg-*Rag1^tm1Mom^ Il2rg^tm1Wjl^/*SzJ), OT-I (C57BL/6-Tg (TcraTcrb)1100Mjb/J), OT-II (B6. Cg-Tg(TcraTcrb)425Cbn/J), B6 CD45.1 (B6.SJL-*Ptprc^a^ Pepc^b^*/BoyJ), male and female mice were procured from the Jackson Laboratory (Marine, USA) and bred in the experimental animal facility (EAF), National Centre for Cell Science (NCCS), Pune, India. All experimental procedures were approved by the Institutional Animal Ethical Committee (IAEC) (Id: EAF/2017/B-256 and EAF/2022/B-441). Six-eight weeks old and age and sex-matched C57BL/6 mice were given subcutaneous injections of MC38, MC38-cOVA, or MC38-HTR_2B_^−/−^ cells (0.5 × 10^6^ cells/mouse in 50 µl PBS). BALB/c mice were given 4T1-Luc2 cells (0.5 X10^6^ cells/mouse) in the mammary fat pad. When mice developed a palpable tumor, treated with intraperitoneal injection of HTR_2B_ agonist (BW-723C86; 4 mg/kg/day), HTR_2B_ antagonist (SB-215505 or RS-127445; 4 mg/kg/day)(*69, 70*), SSRI (Sertraline; 15 mg/kg/twice per week)(*71*), HTR_7_ antagonist (SB-269970; 2.5 mg/kg/day)(*72*), pan-HTR_2_ antagonist (Ketanserin; 2 mg/kg/day), HTR_2A_ antagonist (Ritanserin; 4 mg/kg/day). The largest (height) and the smallest diameter (width) were measured using a digital Vernier caliper, and tumor volume was calculated as described earlier (*73*). Tumor volume = Height × Width^2^ /2.

### Generation of MC38-cOVA and MC38-HTR_2B_ knockout cell lines

To develop the MC38 cells expressing cytoplasmic ovalbumin, MC38 cells were transfected with pCL-Neo-cOVA plasmid [Addgene plasmid # 25097](*29*) using lipofectamine 3000 transfection reagent (ThermoFisher Scientific) according to manufacturer’s guidelines. After 72 hours of transfection, cells were selected using G418 antibiotic (800 μg/ml). Expression of MHC I bound cOVA was assayed after staining with APC labeled anti-mouse H-2K^b^ bound to SIINFEKL antibody (BioLegend) using a flow cytometer or fluorescence microscope.

MC38 cells were transfected with 3 μg of HTR_2B_ CRISPR/Cas9 KO plasmid (Santa Cruz Biotechnology; sc-420994) using lipofectamine 3000 reagents as per manufacturer’s guidelines. After 96 hours, cells were observed for eGFP expression using a flow cytometer, and single cells were sorted in a 96-well plate using a flow cytometer sorter (Moflo Astrios, Beckman Coulter). The eGFP^+^ wells were then selected and expanded. The expression of HTR_2B_ RNA was monitored by real-time qRT-PCR, and the protein was measured by flow cytometry using an anti-HTR_2B_ antibody (Santa Cruz Biotechnology; sc-376878). Mouse IgG1 k isotype antibody was used as the control for flow cytometry.

### Adoptive transfer experiments

MC38 tumors were subcutaneously injected into the C57BL/6 mice. Once palpable tumor developed, mice were treated with an HTR_2B_-antagonist (RS-127445) or sham-treated. On the 10^th^ day, tumor-infiltrating CD8 T cells were magnetically enriched and isolated using flow cytometric sorting. CD8 T cells (1 × 10^6^ cells/mouse) were adoptively transferred into NRG mice (receiving the s.c. injection of MC38 or MC38-cOVA cells). These mice were either treated with an HTR_2B_-antagonist (RS-127445) or sham-treated and on the 10^th^ day, intratumoral CD8 T cells were purified using a flow cytometry cell sorter. Purified CD8 T cells were labeled with Cell-trace™ violet dye (CTV; 10 nM) and were adoptively transferred into congenic B6 CD45.1 mice (1 × 10^6^ cells/mouse) having seven days old MC38-cOVA tumors. The proliferation of CTV-labeled OT-I CD8 T cells was monitored using flow cytometry.

### Depletion of CD8 T cells

Mice were injected with anti-CD8α mAb (clone-YTS 169.4; 100 μg/injection) intravenously every 4^th^ day. Depletion of CD8 T cells in the spleen or lymph nodes was analyzed using flow cytometry.

### Ova-specific CD4 and CD8 T cell culture

C57BL/6 mice were inoculated with MC38-cOVA cell lines, and after the tumor was palpable, mice were given an intraperitoneal injection of HTR_2B_ antagonist (RS-127445; 4 mg/kg/day) for ten days or sham-treated. Mice were euthanized, and single-cell suspensions from tumor-draining lymph nodes (tdLN) were prepared. Cells were then labeled with CFSE (5 nM). CFSE-labeled lymph node cells (5 × 10^4^ of those cells/ well) were cultured in complete RPMI-1640 medium supplemented with ovalbumin protein (50 mg/ml) in U-bottomed 96 well plates at 37°C for four days. Cells were harvested and stained with an anti-mouse CD8 antibody. The ova-specific CD8 T cell proliferation was monitored by CFSE dye dilution using flow cytometry.

### Hematoxylin and Eosin (H&E) staining of FFPE tissue

Tumor tissue samples were excised and fixed in 10% neutral buffered formalin for 48 hours. Tissue samples were dehydrated by treatment with graded concentrations of ethanol (50% for 1 hour twice > 70% for 1 hour twice> 90% for 1 hour twice > 100% for 1 hour twice). Tissue samples were twice treated with xylene for 1 hour, then embedded in liquid paraffin (Paraplast; Sigma) in the embedding chamber to generate formalin-fixed paraffin-embedded (FFPE) tissues. 10 μm thick sections were cut using a microtome and fixed on the positively coated slide. For H&E staining, paraffin was removed from tissue sections by immersing the slides in xylene for 10 minutes, followed by rehydration by immersing them in graded concentrations of ethanol (100% for 10 minutes > 70% for 10 minutes> 50% for 1 hour twice > distilled water for 10 minutes). Then, the tissues were immersed in Harris’s hematoxylin for 10 minutes and matured in tap water for 10 minutes. Then, the tissues were immersed in 1% acid alcohol for destaining, followed by saturated lithium carbonate solution for bluing the dye. Then, the tissues were immersed in 95% ethanol once and then incubated with eosin for 30 seconds. Then, the tissues were again dehydrated using ethanol, followed by cleaning with xylene. Then, images of the mounted tissues were acquired using brightfield microscopy in a Leica DMI6000 fluorescence microscope (Leica Microsystems; Wetzlar, Germany).

### Immunohistochemical staining

Tissue samples were snap-frozen in OCT tissue freezing media (Fisher Scientific) and stored at −80°C. Tissue sections (7 μm thick) were fixed in chilled acetone for 10 mins, air dried, washed with ice-cold PBST (0.1 mM PBS, 0.05% Tween-20), and blocked with 10% horse serum (Jackson ImmunoResearch, West Grove, P.A.) at room temperature (RT) for 1 hour. Sections were washed with PBST and incubated with fluorochrome-tagged or purified primary antibody (1:100 dilutions) in 1% horse serum for four hours at room temperature or 4°C overnight. Sections are washed twice with PBST and stained with secondary antibody (1:800 dilutions) at RT for 1 hour. Then, sections were washed thrice with PBST, fixed with 1% paraformaldehyde, and mounted with DAPI-containing aqueous mounting media (Electron Microscopy Sciences, Hatfield, PA). The images were captured using an Olympus FV3000 confocal microscope (Waltham, Massachusetts) or Leica DMI6000 fluorescence microscope (Leica microsystems; Wetzlar, Germany), and the images were analyzed using Cellsens (Olympus; Waltham, Massachusetts) or Leica LasX software (Leica microsystems; Wetzlar, Germany) or ImageJ software.

### RNA isolation and semi-quantitative PCR

Total RNA was isolated from purified cells using TRIZOL reagent (Invitrogen). RNA was measured, and DNA contamination was removed using DNase I. cDNA was prepared using an Omniscript RT kit (Qiagen), and qRT-PCR was performed using a specific forward and reverse primer mix and a universal SYBR green reverse transcriptase master mix (Biorad; catalog # 1725122) in CFX96 Real-Time PCR system (Biorad).

### Preparation of single-cell suspension and cell staining for flow cytometry

Tumors were excised, manually disrupted into small pieces using fine forceps and scissors, and resuspended in serum-free Hank’s balanced salt solution (HBSS) containing collagenase type I (0.1 mg/ml), collagenase IV (0.1 mg/ml), collagenase D (0.1 mg/ml), hyaluronidase (0.06 mg/ml), DNase I (0.02 mg/ml), Dispase I (0.25 mg/ml) and soybean trypsin inhibitor (0.1 mg/ml) and incubated at 37°C in a shaker incubator for 30 minutes. The tissue pieces were then resuspended in complete RPMI media, and single-cell suspension was prepared by passing through a 70 μM pore-size cell strainer. Then, cells were washed with PBS and stained with fluorochrome-tagged primary antibodies. Single-cell suspension of spleen and lymph nodes was prepared by mechanical disruption of the tissues, and the cell suspension was passed through a 70 μm pore size cell strainer. RBCs were removed using an ACK lysis buffer and were washed with a complete RPMI medium. Cells were incubated with fluorochrome-tagged primary antibodies on ice for 1 hour, followed by one wash with PBS, and cells were either fixed with 1 % paraformaldehyde or proceeded for intracellular staining. For intracellular staining, cells were fixed with 1X FoxP3 fixation buffer (BioLegend) on ice for 45 minutes, followed by washing with 1X permeabilization buffer (BioLegend). Then, the cells were incubated with a permeabilization buffer for 10 minutes on ice. Cells were then incubated with fluorochrome-tagged specific antibodies in permeabilization buffer for 1 hour, washed with PBS, and fixed with 0.5% paraformaldehyde. Finally, the cells were acquired in flow cytometry (Canto II, BD Biosciences; or Cytek Aurora, Cytek Biosciences), and the data were analyzed using FlowJo, FCS express, or OMIQ software.

### Intracellular cytokine staining for flow cytometry

For intracellular cytokine staining, a single-cell suspension was prepared from lymph nodes or tumors. Cells were cultured in Phorbol 12-myristate 13-acetate (PMA; 50 ng/mL) and Ionomycin (500 ng/mL) at 37℃ in a 5% CO_2_ incubator. For most of the cytokines, cells were cultured for 6 hours, but for IL-4 and IL-10, cells were cultured for 10 hours. After 90 minutes of culture initiation, 1X brefeldin A (BioLegend) was added to each well to block the secretion of the cytokine. Cells were harvested and first stained for surface markers and then fixed using 1X cytoplasm fix buffer (Biolegend) on ice for 45 mins, followed by 1X Cytofast permeabilization on ice for 5 mins. Cells were incubated in the antibody cocktails against the cytokines in permeabilization buffer for 30 mins, washed in permeabilization buffer once, and followed by one wash with PBS. The cells were acquired in a flow cytometer (Canto II or Cytek Aurora), and data were analyzed using FlowJo, FCS express, or OMIQ software.

### Tetramer staining

From the single cell suspensions prepared for flow cytometry staining, the sample was stained with fixable live-dead staining (Cytek) for 20 minutes in PBS on ice. Cells were washed once with cell staining buffer and incubated in complete RPMI media with OT-I tetramer (1:100 dilution) at 37°C for 30 minutes or with OT-II tetramer (1:10 dilution) at 37°C for 2 hours. Cells were washed once with a cell staining buffer and stained for other surfaces and intracellular markers.

### Analysis of multi-dimensional data of FACS

High-dimensional data from flow cytometry was analyzed using OMIQ cloud-based software. The data were cleaned by gating the singlets and the cellular populations from FSC and SSC plots. The live cells and subsequent cellular populations were identified. Subsampling was performed on the cellular population, and an equal representative cell number from each sample was taken. Dimensionality reduction was performed using the UMAP (Uniform Manifold Approximation and Projection) algorithm using the basic parameters such as Neighbors-15, Minimum distance-0.8, components-2, metric-euclidean, learning rate-1, epochs-200, random seeds-42, embedding initialization-spectral. Unsupervised clustering was performed on the UMAP using the FlowSom algorithm using basing settings, and consensus meta-clustering was performed. The cellular identity of the meta clusters was identified from the clustered heatmap. The results were visualized with a density or colored continuous scatter plot.

### ELISA

Universal competitive serotonin ELISA kit (Novus biological; Catalog # NBP2-68127), Mouse PD-L1 DuoSet ELISA kit (R&D System; Catalog# DY1019-05), mouse IL-10 ELISA MAX kit (BioLegend; catalog# 431414), and Mouse TNF-α ELISA MAX kit (BioLegend; catalog# 430904) were used to perform ELISA according to the manufacturer’s guidelines.

### Chromogenic immunohistochemistry

The human colon adenocarcinoma tissue microarray (Novus Biological, Catalog # NBP2-78088) was stained with the antibodies and detected using chromogenic reagent DAB. Briefly, the slides were pre-warmed at 60°C for one hour before staining and then deparaffinized using xylene and then dehydrated using sequential treatment of molecular grade ethyl alcohol of graded concentrations (50%>70%>90%.100%). Then, antigen retrieval was performed using antigen-retrieval buffer (BioLegend; catalog # 928001; pH 8.0) at sub-boiling temperature (90 °C) for 15 minutes. Then, the intrinsic peroxidase of the tissue was quenched by Peroxidase blocking reagent (BioLegend; catalog # 927402), followed by blocking and permeabilization using 10% goat serum in PBST (0.1 mM PBS; 0.5% Triton-X). The sections were further incubated with primary antibodies such as rabbit polyclonal anti-human HTR_2B_ antibody (1:100 dilution) or rabbit polyclonal anti-human TPH1 (1:100 dilution) at 4°C overnight. The slides were washed twice with PBST (1X PBS, 0.05% Tween 20) and incubated with a goat anti-rabbit HRP-tagged secondary IgG (1:200) at room temperature for 1 hour. Sections were probed with DAB-H_2_O_2_ reagent and counterstained with hematoxylin, and images were acquired using bright field microscopy (Leica DMI6000 fluorescence microscope, Leica microsystems; Wetzlar, Germany).

### 4T1-Luc2 lung metastasis model

4T1-Luc2 cells were cultured in RPMI-1640 medium at a 60-70% confluency. Cells were harvested, and 5 × 10^5^ cells/mice were injected into female BALB/c mice intravenously (150 μl) through the tail vein. Mice were treated with either HTR_2B_ antagonist (RS-127445; 4 mg/kg/day) for at least 10 days or left untreated. Then, the extent of lung metastasis was visualized using the IVIS® spectrum *in vivo* imaging system (PerkinElmer) and analyzed using Living image® software (PerkinElmer) after injecting the mice with D-luciferin (150 mg/kg; PerkinElmer) in PBS. Mice were further euthanized, and the number of visible nodules on the lung surfaces was monitored.

### TCGA survival data analysis

The correlation between serotonin receptor expression pattern and patient overall survival (OS) with colon adenocarcinoma (COAD) was analyzed through the oncolnc.org website using The Cancer Genome Atlas (TCGA) database (*74*). The survival status of COAD subjects from TCGA databases showing the highest expression (top 25% segment; n=110) of serotonin receptor subtypes and COAD subjects showing the lowest expression (bottom 25% segment; n=110) of serotonin receptor subtypes were correlated, and the Kaplan-Mayer plot for every serotonin receptor subtype was plotted.

### Tumor immune dysfunction and exclusion (TIDE) computational method

TIDE analysis was conducted utilizing the platform at http://tide.dfci.harvard.edu/. This method was employed to investigate the relationship between tumor-infiltrating CD8^+^ cytotoxic T lymphocytes (CTLs) and overall survival in the context of HTR_2B_ gene expression. Tumor samples were stratified into two groups based on HTR_2B_ expression: HTR_2B_-high (samples with expression levels greater than one standard deviation above the mean) and HTR_2B_-low (remaining samples). The association between CTL levels and survival outcomes was then analyzed for each group. CTL functionality was quantified by assessing the mean expression levels of CD8A, CD8B, GZMA, GZMB, and PRF1. Survival plots were generated for two subgroups: “CTL-High,” characterized by above-average CTL values across all samples, and “CTL-Low,” representing samples with below-average CTL levels. A T cell dysfunction z score was computed for each cohort, evaluating the correlation between HTR_2B_ expression and the efficacy of CTL infiltration in promoting survival. A positive z score reflects a negative correlation between HTR_2B_ expression and the beneficial impact of tumor-infiltrating CTLs on patient survival.

### Reanalysis of TCGA-COAD RNA sequencing datasets

Based on HTR_2B_ expression, 30 samples from TCGA-COAD datasets are downloaded from the NCBI GDC portal (https://portal.gdc.cancer.gov/repository). Among them, 15 samples are characterized as HTR_2B_-High, expressing the highest HTR_2B,_ and the other 15 samples are characterized as HTR_2B_ low (patient details are given in **Supplementary Table 2**). From the RNA-seq datasets, the genes with very low read counts were removed, and the rest of the data was analyzed using R studio [RStudio Team (2021). RStudio: Integrated Development Environment for R. RStudio, PBC, Boston, MA URL http://www.rstudio.com)]. The differential expression analysis of different genes was performed using the DESeq2 package in R (*75*). The volcano plots were generated using the Enhanced Volcano package in R studio— the cluster profiler package performed gene ontology analysis (*76*). The gene set enrichment analysis (GSEA) was performed using the GSEA software from Broad Institute (*77*). For angiogenesis and Epithelial-mesenchymal transition, the GSEA was performed on gene sets from the Molecular signature database (MSigDB) like the Hallmark gene sets (*78*), GOBP gene sets for serotonin receptor signaling pathway (GO:0007210) and serotonin transport (GO:0006837), and KEGG pathway gene sets for T cell receptor signaling (hsa04660), cytokine-to-cytokine receptor interaction (hsa04060), chemokine signaling pathway (hsa04662, and TGFβ signaling pathway (hsa04350) were analyzed. Apart from these gene sets, the gene sets associated with T cell effector function **(Supplementary Table 3)** and T cell regulatory functions **(Supplementary Table 3)** were also used to perform GSEA. The Heat map was generated using Morpheus (https://software.broadinstitute.org/morpheus) software.

### Statistical analysis

Unpaired two-tailed Student’s t-test was used to compare two independent groups. One-way ANOVA with Tukey’s multiple comparison test was used to compare the effects of one variable. Two-way ANOVA with Sidak’s multiple comparison test was used to compare the impacts of more than one variable on two or more independent experimental groups. A p-value of less than 0.05 was considered statistically significant. Unless otherwise indicated, all the statistical data comparisons are represented as mean ± Standard Error of Mean (SEM). All statistical analyses were performed using GraphPad Prism 10 software (GraphPad Software, San Diego, CA).

## Author’s contribution

GL conceived the idea, designed the experiments, and arranged the resources and funding. SK designed and performed the experiments and analyzed the data. GL and SK wrote and finalized the manuscript.

## Acknowledgment

GL received a Swarna Jayanti Fellowship (DST/SJF/LSA-01/2017-2018) from the Department of Science and Technology, a TATA Innovation Fellowship (HRD-16012/5/2024) from the Department of Biotechnology, and a research grant from the Science and Engineering Research Board (CRG/2022/007108), Ministry of Science and Technology, Government of India. SK received a Senior Research Fellowship from the Council of Scientific and Industrial Research, Government of India.

## References

1. R. M. Sobel, D. Markov, The impact of anxiety and mood disorders on physical disease: the worried not-so-well. Curr Psychiatry Rep 7, 206–212 (2005).

2. S. H. Lin, L. T. Lee, Y. K. Yang, Serotonin and mental disorders: a concise review on molecular neuroimaging evidence. Clin Psychopharmacol Neurosci 12, 196–202 (2014).

3. D. J. Walther, J. U. Peter, S. Bashammakh, H. Hörtnagl, M. Voits, H. Fink, M. Bader, Synthesis of serotonin by a second tryptophan hydroxylase isoform. Science 299, 1078197 (2003).

4. N. Herr, C. Bode, D. Duerschmied, The Effects of Serotonin in Immune Cells. Frontiers in Cardiovascular Medicine 4, (2017).

5. J. Masson, M. B. Emerit, M. Hamon, M. Darmon, Serotonergic signaling: multiple effectors and pleiotropic effects. Wiley Interdisciplinary Reviews: Membrane Transport and Signaling 1, 685–713 (2012).

6. M. M. Rapport, A. A. Green, I. H. Page, Serum vasoconstrictor, serotonin; isolation and characterization. J Biol Chem 176, 1243–1251 (1948).

7. A. Zamani, Z. Qu, Serotonin activates angiogenic phosphorylation signaling in human endothelial cells. FEBS Lett 586, 2360–2365 (2012).

8. V. K. Yadav, S. Balaji, P. S. Suresh, X. S. Liu, X. Lu, Z. Li, X. E. Guo, J. J. Mann, A. K. Balapure, M. D. Gershon, R. Medhamurthy, M. Vidal, G. Karsenty, P. Ducy, Pharmacological inhibition of gut-derived serotonin synthesis is a potential bone anabolic treatment for osteoporosis. Nat Med 16, 308–312 (2010).

9. M. Berger, J. A. Gray, B. L. Roth, The expanded biology of serotonin. Annu Rev Med 60, 355–366 (2009).

10. A. M. Martin, R. L. Young, L. Leong, G. B. Rogers, N. J. Spencer, C. F. Jessup, D. J. Keating, The Diverse Metabolic Roles of Peripheral Serotonin. Endocrinology 158, 1049–1063 (2017).

11. S. Karmakar, G. Lal, Role of serotonin receptor signaling in cancer cells and anti-tumor immunity. Theranostics 11, 5296–5312 (2021).

12. S. Karmakar, G. Lal, in Neuroprotection: Method and Protocols, S. K. Ray, Ed. (Springer US, New York, NY, 2024), pp. 181–207.

13. M. de las Casas-Engel, A. Domínguez-Soto, E. Sierra-Filardi, R. Bragado, C. Nieto, A. Puig-Kroger, R. Samaniego, M. Loza, M. T. Corcuera, F. Gómez-Aguado, M. Bustos, P. Sánchez-Mateos, A. L. Corbí, Serotonin Skews Human Macrophage Polarization through HTR_2B_ and HTR_7_. The Journal of Immunology 190, 2301–2310 (2013).

14. A. Szabo, P. Gogolak, G. Koncz, Z. Foldvari, K. Pazmandi, N. Miltner, S. Poliska, A. Bacsi, S. Djurovic, E. Rajnavolgyi, Immunomodulatory capacity of the serotonin receptor 5-HT2B in a subset of human dendritic cells. Sci Rep 8, 018–20173 (2018).

15. M. Idzko, E. Panther, C. Stratz, T. Müller, H. Bayer, G. Zissel, T. Dürk, S. Sorichter, F. Di Virgilio, M. Geissler, B. Fiebich, Y. Herouy, P. Elsner, J. Norgauer, D. Ferrari, The serotoninergic receptors of human dendritic cells: identification and coupling to cytokine release. Journal of immunology (Baltimore, Md. : 1950) 172, 6011–6019 (2004).

16. L. Zhang, S. Yang, X. Chen, S. Stauffer, F. Yu, S. M. Lele, K. Fu, K. Datta, N. Palermo, Y. Chen, J. Dong, The hippo pathway effector YAP regulates motility, invasion, and castration-resistant growth of prostate cancer cells. Mol Cell Biol 35, 1350–1362 (2015).

17. B. Shu, M. Zhai, X. Miao, C. He, C. Deng, Y. Fang, M. Luo, L. Liu, S. Liu, Serotonin and YAP/VGLL4 Balance Correlated with Progression and Poor Prognosis of Hepatocellular Carcinoma. Sci Rep 8, 9739 (2018).

18. S. Liu, R. Miao, M. Zhai, Q. Pang, Y. Deng, S. Liu, K. Qu, C. Liu, J. Zhang, Effects and related mechanisms of serotonin on malignant biological behavior of hepatocellular carcinoma via regulation of Yap. Oncotarget 8, 47412–47424 (2017).

19. E. J. Siddiqui, M. Shabbir, D. P. Mikhailidis, C. S. Thompson, F. H. Mumtaz, The role of serotonin (5-hydroxytryptamine1A and 1B) receptors in prostate cancer cell proliferation. J Urol 176, 1648–1653 (2006).

20. R. Ataee, S. Ajdary, M. Zarrindast, M. Rezayat, M. R. Hayatbakhsh, Anti-mitogenic and apoptotic effects of 5-HT1B receptor antagonist on HT29 colorectal cancer cell line. J Cancer Res Clin Oncol 136, 1461–1469 (2010).

21. M. W. Rosenbaum, J. R. Bledsoe, V. Morales-Oyarvide, T. G. Huynh, M. Mino-Kenudson, PD-L1 expression in colorectal cancer is associated with microsatellite instability, BRAF mutation, medullary morphology and cytotoxic tumor-infiltrating lymphocytes. Modern Pathology 29, 1104–1112 (2016).

22. X. Chen, L.-J. Chen, X.-F. Peng, L. Deng, Y. Wang, J.-J. Li, D.-L. Guo, X.-H. Niu, Anti-PD-1/PD-L1 therapy for colorectal cancer: Clinical implications and future considerations. Translational Oncology 40, 101851 (2024).

23. T. J. Miller, C. C. Anyaegbu, T. F. Lee-Pullen, L. J. Spalding, C. F. Platell, M. J. McCoy, PD-L1+ dendritic cells in the tumor microenvironment correlate with good prognosis and CD8+ T cell infiltration in colon cancer. Cancer science 112, 1173–1183 (2021).

24. A. M. Valentini, F. Di Pinto, F. Cariola, V. Guerra, G. Giannelli, M. L. Caruso, M. Pirrelli, PD-L1 expression in colorectal cancer defines three subsets of tumor immune microenvironments. Oncotarget 9, 8584–8596 (2018).

25. T. Shan, S. Chen, T. Wu, Y. Yang, S. Li, X. Chen, PD-L1 expression in colon cancer and its relationship with clinical prognosis. International journal of clinical and experimental pathology 12, 1764–1769 (2019).

26. N. Yaghoubi, A. Soltani, K. Ghazvini, S. M. Hassanian, S. I. Hashemy, PD-1/ PD-L1 blockade as a novel treatment for colorectal cancer. Biomedicine & Pharmacotherapy 110, 312–318 (2019).

27. J. L. Benci, L. R. Johnson, R. Choa, Y. Xu, J. Qiu, Z. Zhou, B. Xu, D. Ye, K. L. Nathanson, C. H. June, E. J. Wherry, N. R. Zhang, H. Ishwaran, M. D. Hellmann, J. D. Wolchok, T. Kambayashi, A. J. Minn, Opposing Functions of Interferon Coordinate Adaptive and Innate Immune Responses to Cancer Immune Checkpoint Blockade. Cell 178, 933–948.e914 (2019).

28. M. Philip, A. Schietinger, CD8(+) T cell differentiation and dysfunction in cancer. Nat Rev Immunol 22, 209–223 (2022).

29. J. Yang, N. S. Sanderson, K. Wawrowsky, M. Puntel, M. G. Castro, P. R. Lowenstein, Kupfer-type immunological synapse characteristics do not predict anti-brain tumor cytolytic T-cell function in vivo. Proc Natl Acad Sci U S A 107, 4716–4721 (2010).

30. S. H. Karandikar, J. Sidney, A. Sette, M. J. Selby, A. J. Korman, P. K. Srivastava, New epitopes in ovalbumin provide insights for cancer neoepitopes. JCI insight 5, (2019).

31. R. Wannemacher, A. Reiß, K. Rohn, F. Lühder, A. Flügel, W. Baumgärtner, K. Hülskötter, Ovalbumin-specific CD4+ and CD8+ T cells contribute to different susceptibility for Theiler’s murine encephalomyelitis virus persistence. 14, (2023).

32. S. R. Clarke, M. Barnden, C. Kurts, F. R. Carbone, J. F. Miller, W. R. Heath, Characterization of the ovalbumin-specific TCR transgenic line OT-I: MHC elements for positive and negative selection. Immunology and cell biology 78, 110–117 (2000).

33. I. Puzanov, A. Diab, K. Abdallah, C. O. Bingham, C. Brogdon, R. Dadu, L. Hamad, S. Kim, M. E. Lacouture, N. R. LeBoeuf, D. Lenihan, C. Onofrei, V. Shannon, R. Sharma, A. W. Silk, D. Skondra, M. E. Suarez-Almazor, Y. Wang, K. Wiley, H. L. Kaufman, M. S. Ernstoff, o. b. o. t. S. f. I. o. C. T. M. W. Group, J. Anderson, D. Arrindell, S. Andrews, J. Ballesteros, J. Boyer, D. Chen, D. Chonzi, I. Cotarla, R. Cunha, M. Davies, M. Dawson, A. Dicker, L. Eifler, A. Ferguson, C. Ferlini, S. Frankel, W. Go, C. Gochett, J. Goldberg, P. Goncalves, T. Goswami, N. Gregory, J. L. Gulley, V. Hayreh, N. Helie, W. Holmes, J.-Y. Hsu, R. Ibrahim, C. Larocca, K. Lehman, S. Ley-Acosta, O. Lambotte, J. Luke, J. McClure, E. Michelon, M. Nakamura, K. Patel, B. Piperdi, Z. Rasheed, D. Reshef, J. Riemer, C. Robert, M. Sarkeshik, A. Saylors, J. Schreiber, K. Shafer-Weaver, W. Sharfman, E. Sharon, R. Sherry, C. Simonson, C. Thomas, J. A. Thompson, E. Trehu, D. Tresnan, M. Turner, D. Wariabharaj, I. Waxman, L. Wood, L. Zhang, P. Zheng, Managing toxicities associated with immune checkpoint inhibitors: consensus recommendations from the Society for Immunotherapy of Cancer (SITC) Toxicity Management Working Group. 5, 95 (2017).

34. J. A. Marin-Acevedo, R. M. Chirila, R. S. Dronca, Immune Checkpoint Inhibitor Toxicities. Mayo Clinic proceedings 94, 1321–1329 (2019).

35. X. Kong, L. Chen, Z. Su, R. J. Sullivan, S. M. Blum, Z. Qi, Y. Liu, Y. Huo, Y. Fang, L. Zhang, J. Gao, J. Wang, Toxicities associated with immune checkpoint inhibitors: a systematic study. International journal of surgery 109, 1753–1768 (2023).

36. N. Riaz, J. J. Havel, V. Makarov, A. Desrichard, W. J. Urba, J. S. Sims, F. S. Hodi, S. Martín-Algarra, R. Mandal, W. H. Sharfman, S. Bhatia, W. J. Hwu, T. F. Gajewski, C. L. Slingluff, Jr., D. Chowell, S. M. Kendall, H. Chang, R. Shah, F. Kuo, L. G. T. Morris, J. W. Sidhom, J. P. Schneck, C. E. Horak, N. Weinhold, T. A. Chan, Tumor and Microenvironment Evolution during Immunotherapy with Nivolumab. Cell 171, 934–949.e916 (2017).

37. P. Comella, R. Casaretti, C. Sandomenico, A. Avallone, L. Franco, Role of oxaliplatin in the treatment of colorectal cancer. Ther Clin Risk Manag 5, 229–238 (2009).

38. S. Vodenkova, T. Buchler, K. Cervena, V. Veskrnova, P. Vodicka, V. Vymetalkova, 5-fluorouracil and other fluoropyrimidines in colorectal cancer: Past, present and future. Pharmacol Ther 206, 107447 (2020).

39. S. Kawai, N. Takeshima, Y. Hayasaka, A. Notsu, M. Yamazaki, T. Kawabata, K. Yamazaki, K. Mori, H. Yasui, Comparison of irinotecan and oxaliplatin as the first-line therapies for metastatic colorectal cancer: a meta-analysis. BMC Cancer 21, 116 (2021).

40. T. Ling, Z. Dai, H. Wang, T. T. Kien, R. Cui, T. Yu, J. Chen, Serotonylation in tumor-associated fibroblasts contributes to the tumor-promoting roles of serotonin in colorectal cancer. Cancer Letters 600, 217150 (2024).

41. C. G. Nebigil, P. Hickel, N. Messaddeq, J.-L. Vonesch, M. P. Douchet, L. Monassier, K. György, R. Matz, R. Andriantsitohaina, P. Manivet, J.-M. Launay, L. Maroteaux, Ablation of Serotonin 5-HT2B Receptors in Mice Leads to Abnormal Cardiac Structure and Function. Circulation 103, 2973–2979 (2001).

42. C. G. Nebigil, D. S. Choi, A. Dierich, P. Hickel, M. Le Meur, N. Messaddeq, J. M. Launay, L. Maroteaux, Serotonin 2B receptor is required for heart development. Proc Natl Acad Sci U S A 97, 9508–9513 (2000).

43. C. Burgess, V. Cornelius, S. Love, J. Graham, M. Richards, A. Ramirez, Depression and anxiety in women with early breast cancer: five year observational cohort study. Bmj 330, 702 (2005).

44. Y. N. Peng, M. L. Huang, C. H. Kao, Prevalence of Depression and Anxiety in Colorectal Cancer Patients: A Literature Review. Int J Environ Res Public Health 16, (2019).

45. F. Mols, D. Schoormans, I. de Hingh, S. Oerlemans, O. Husson, Symptoms of anxiety and depression among colorectal cancer survivors from the population-based, longitudinal PROFILES Registry: Prevalence, predictors, and impact on quality of life. Cancer 124, 2621–2628 (2018).

46. M. E. Renna, M. R. Shrout, A. A. Madison, C. M. Alfano, S. P. Povoski, A. M. Lipari, W. E. Carson, 3rd, W. B. Malarkey, J. K. Kiecolt-Glaser, Depression and anxiety in colorectal cancer patients: Ties to pain, fatigue, and inflammation. Psychooncology 31, 1536–1544 (2022).

47. E. F. Cascade, A. H. Kalali, M. E. Thase, Use of antidepressants: expansion beyond depression and anxiety. Psychiatry (Edgmont) 4, 25–28 (2007).

48. V. Abbing-Karahagopian, C. Huerta, P. C. Souverein, F. de Abajo, H. G. Leufkens, J. Slattery, Y. Alvarez, M. Miret, M. Gil, B. Oliva, U. Hesse, G. Requena, F. de Vries, M. Rottenkolber, S. Schmiedl, R. Reynolds, R. G. Schlienger, M. C. de Groot, O. H. Klungel, T. P. van Staa, L. van Dijk, A. C. Egberts, H. Gardarsdottir, M. L. De Bruin, Antidepressant prescribing in five European countries: application of common definitions to assess the prevalence, clinical observations, and methodological implications. Eur J Clin Pharmacol 70, 849–857 (2014).

49. B. Boursi, I. Lurie, K. Haynes, R. Mamtani, Y. X. Yang, Chronic therapy with selective serotonin reuptake inhibitors and survival in newly diagnosed cancer patients. Eur J Cancer Care (Engl) 27, (2018).

50. J. Busby, K. Mills, S. D. Zhang, F. G. Liberante, C. R. Cardwell, Selective serotonin reuptake inhibitor use and breast cancer survival: a population-based cohort study. Breast Cancer Res 20, 4 (2018).

51. A. Fischer, H. S. Rennert, G. Rennert, Selective serotonin reuptake inhibitors associated with increased mortality risk in breast cancer patients in Northern Israel. Int J Epidemiol 51, 807–816 (2022).

52. D. Hirano, Y. Urabe, S. Tanaka, K. Nakamura, Y. Ninomiya, R. Yuge, R. Hayashi, S. Oka, Y. Kitadai, F. Shimamoto, K. Arihiro, K. Chayama, Early-stage serrated adenocarcinomas are divided into several molecularly distinct subtypes. PLoS ONE 14, e0211477 (2019).

53. T. Li, L. Wei, X. Zhang, B. Fu, Y. Zhou, M. Yang, M. Cao, Y. Chen, Y. Tan, Y. Shi, L. Wu, C. Xuan, Q. Du, R. Hu, Serotonin Receptor HTR2B Facilitates Colorectal Cancer Metastasis via CREB1-ZEB1 Axis-Mediated Epithelial-Mesenchymal Transition. Mol Cancer Res 22, 538–554 (2024).

54. T. Ling, Z. Dai, H. Wang, T. T. Kien, R. Cui, T. Yu, J. Chen, Serotonylation in tumor-associated fibroblasts contributes to the tumor-promoting roles of serotonin in colorectal cancer. Cancer letters 600, 217150 (2024).

55. J. E. Ghia, N. Li, H. Wang, M. Collins, Y. Deng, R. T. El-Sharkawy, F. Cote, J. Mallet, W. I. Khan, Serotonin has a key role in pathogenesis of experimental colitis. Gastroenterology 137, 1649–1660 (2009).

56. L. Mao, F. Xin, J. Ren, S. Xu, H. Huang, X. Zha, X. Wen, G. Gu, G. Yang, Y. Cheng, C. Zhang, W. Wang, X. Liu, 5-HT2B-mediated serotonin activation in enterocytes suppresses colitis-associated cancer initiation and promotes cancer progression. Theranostics 12, 3928–3945 (2022).

57. S. J. Koh, J. M. Kim, I. K. Kim, N. Kim, H. C. Jung, I. S. Song, J. S. Kim, Fluoxetine inhibits NF-kappaB signaling in intestinal epithelial cells and ameliorates experimental colitis and colitis-associated colon cancer in mice. American journal of physiology 301, G9–19 (2011).

58. J. L. Montalvo-Ortiz, H. Zhou, I. D’Andrea, L. Maroteaux, A. Lori, A. Smith, K. J. Ressler, Y. Z. Nunez, L. A. Farrer, H. Zhao, H. R. Kranzler, J. Gelernter, Translational studies support a role for serotonin 2B receptor (HTR2B) gene in aggression-related cannabis response. Mol Psychiatry 23, 2277–2286 (2018).

59. S. L. Diaz, S. Doly, N. Narboux-Neme, S. Fernandez, P. Mazot, S. M. Banas, K. Boutourlinsky, I. Moutkine, A. Belmer, A. Roumier, L. Maroteaux, 5-HT(2B) receptors are required for serotonin-selective antidepressant actions. Mol Psychiatry 17, 154–163 (2012).

60. P. M. Pitychoutis, A. Belmer, I. Moutkine, J. Adrien, L. Maroteaux, Mice Lacking the Serotonin Htr2B Receptor Gene Present an Antipsychotic-Sensitive Schizophrenic-Like Phenotype. Neuropsychopharmacology 40, 2764–2773 (2015).

61. C. Bechade, I. D’Andrea, F. Etienne, F. Verdonk, I. Moutkine, S. M. Banas, M. Kolodziejczak, S. L. Diaz, C. N. Parkhurst, W. B. Gan, L. Maroteaux, A. Roumier, The serotonin 2B receptor is required in neonatal microglia to limit neuroinflammation and sickness behavior in adulthood. Glia 69, 638–654 (2021).

62. Y. Y. Janjigian, K. Shitara, M. Moehler, M. Garrido, P. Salman, L. Shen, L. Wyrwicz, K. Yamaguchi, T. Skoczylas, A. Campos Bragagnoli, T. Liu, M. Schenker, P. Yanez, M. Tehfe, R. Kowalyszyn, M. V. Karamouzis, R. Bruges, T. Zander, R. Pazo-Cid, E. Hitre, K. Feeney, J. M. Cleary, V. Poulart, D. Cullen, M. Lei, H. Xiao, K. Kondo, M. Li, J. A. Ajani, First-line nivolumab plus chemotherapy versus chemotherapy alone for advanced gastric, gastro-oesophageal junction, and oesophageal adenocarcinoma (CheckMate 649): a randomised, open-label, phase 3 trial. Lancet (London, England) 398, 27–40 (2021).

63. U. Sahin, P. Oehm, E. Derhovanessian, R. A. Jabulowsky, M. Vormehr, M. Gold, D. Maurus, D. Schwarck-Kokarakis, A. N. Kuhn, T. Omokoko, L. M. Kranz, M. Diken, S. Kreiter, H. Haas, S. Attig, R. Rae, K. Cuk, A. Kemmer-Brück, A. Breitkreuz, C. Tolliver, J. Caspar, J. Quinkhardt, L. Hebich, M. Stein, A. Hohberger, I. Vogler, I. Liebig, S. Renken, J. Sikorski, M. Leierer, V. Müller, H. Mitzel-Rink, M. Miederer, C. Huber, S. Grabbe, J. Utikal, A. Pinter, R. Kaufmann, J. C. Hassel, C. Loquai, Ö. Türeci, An RNA vaccine drives immunity in checkpoint-inhibitor-treated melanoma. Nature 585, 107–112 (2020).

64. J. Larkin, V. Chiarion-Sileni, R. Gonzalez, J. J. Grob, C. L. Cowey, C. D. Lao, D. Schadendorf, R. Dummer, M. Smylie, P. Rutkowski, P. F. Ferrucci, A. Hill, J. Wagstaff, M. S. Carlino, J. B. Haanen, M. Maio, I. Marquez-Rodas, G. A. McArthur, P. A. Ascierto, G. V. Long, M. K. Callahan, M. A. Postow, K. Grossmann, M. Sznol, B. Dreno, L. Bastholt, A. Yang, L. M. Rollin, C. Horak, F. S. Hodi, J. D. Wolchok, Combined Nivolumab and Ipilimumab or Monotherapy in Untreated Melanoma. The New England journal of medicine 373, 23–34 (2015).

65. H. Borghaei, L. Paz-Ares, L. Horn, D. R. Spigel, M. Steins, N. E. Ready, L. Q. Chow, E. E. Vokes, E. Felip, E. Holgado, F. Barlesi, M. Kohlhäufl, O. Arrieta, M. A. Burgio, J. Fayette, H. Lena, E. Poddubskaya, D. E. Gerber, S. N. Gettinger, C. M. Rudin, N. Rizvi, L. Crinò, G. R. Blumenschein, Jr., S. J. Antonia, C. Dorange, C. T. Harbison, F. Graf Finckenstein, J. R. Brahmer, Nivolumab versus Docetaxel in Advanced Nonsquamous Non-Small-Cell Lung Cancer. The New England journal of medicine 373, 1627–1639 (2015).

66. M. Mollaei, Z. M. Hassan, F. Khorshidi, L. Langroudi, Chemotherapeutic drugs: Cell death- and resistance-related signaling pathways. Are they really as smart as the tumor cells? Transl Oncol 14, 101056 (2021).

67. S. U. Khan, K. Fatima, S. Aisha, F. Malik, Unveiling the mechanisms and challenges of cancer drug resistance. Cell Communication and Signaling 22, 109 (2024).

68. R. W. Robey, K. M. Pluchino, M. D. Hall, A. T. Fojo, S. E. Bates, M. M. Gottesman, Revisiting the role of ABC transporters in multidrug-resistant cancer. Nature Reviews Cancer 18, 452–464 (2018).

69. A. Fabre, J. Marchal-Sommé, S. Marchand-Adam, C. Quesnel, R. Borie, M. Dehoux, C. Ruffié, J. Callebert, J. M. Launay, D. Hénin, P. Soler, B. Crestani, Modulation of bleomycin-induced lung fibrosis by serotonin receptor antagonists in mice. Eur Respir J 32, 426–436 (2008).

70. D. W. Bonhaus, L. A. Flippin, R. J. Greenhouse, S. Jaime, C. Rocha, M. Dawson, K. Van Natta, L. K. Chang, T. Pulido-Rios, A. Webber, E. Leung, R. M. Eglen, G. R. Martin, RS-127445: a selective, high affinity, orally bioavailable 5-HT2B receptor antagonist. Br J Pharmacol 127, 1075–1082 (1999).

71. I. Gil-Ad, A. Zolokov, L. Lomnitski, M. Taler, M. Bar, D. Luria, E. Ram, A. Weizman, Evaluation of the potential anti-cancer activity of the antidepressant sertraline in human colon cancer cell lines and in colorectal cancer-xenografted mice. Int J Oncol 33, 277–286 (2008).

72. J. Sowa, M. Kusek, M. Siwiec, J. E. Sowa, B. Bobula, K. Tokarski, G. Hess, The 5-HT. Psychopharmacology (Berl) 235, 3381–3390 (2018).

73. M. M. Jensen, J. T. Jørgensen, T. Binderup, A. Kjaer, Tumor volume in subcutaneous mouse xenografts measured by microCT is more accurate and reproducible than determined by 18F-FDG-microPET or external caliper. BMC Med Imaging 8, 16 (2008).

74. J. Anaya, OncoLnc: linking TCGA survival data to mRNAs, miRNAs, and lncRNAs. PeerJ Computer Science 2, e67 (2016).

75. M. I. Love, W. Huber, S. Anders, Moderated estimation of fold change and dispersion for RNA-seq data with DESeq2. Genome Biol 15, 550 (2014).

76. T. Wu, E. Hu, S. Xu, M. Chen, P. Guo, Z. Dai, T. Feng, L. Zhou, W. Tang, L. Zhan, X. Fu, S. Liu, X. Bo, G. Yu, clusterProfiler 4.0: A universal enrichment tool for interpreting omics data. Innovation (Camb) 2, 100141 (2021).

77. V. K. Mootha, C. M. Lindgren, K. F. Eriksson, A. Subramanian, S. Sihag, J. Lehar, P. Puigserver, E. Carlsson, M. Ridderstråle, E. Laurila, N. Houstis, M. J. Daly, N. Patterson, J. P. Mesirov, T. R. Golub, P. Tamayo, B. Spiegelman, E. S. Lander, J. N. Hirschhorn, D. Altshuler, L. C. Groop, PGC-1alpha-responsive genes involved in oxidative phosphorylation are coordinately downregulated in human diabetes. Nat Genet 34, 267–273 (2003).

78. A. Liberzon, C. Birger, H. Thorvaldsdóttir, M. Ghandi, J. P. Mesirov, P. Tamayo, The Molecular Signatures Database (MSigDB) hallmark gene set collection. Cell Syst 1, 417–425 (2015).

