## Supplementary material for "Antagonizing the serotonin receptor HTR2B drives antigen-specific cytotoxic T-cell responses and controls colorectal cancer growth": Figures S1-S3

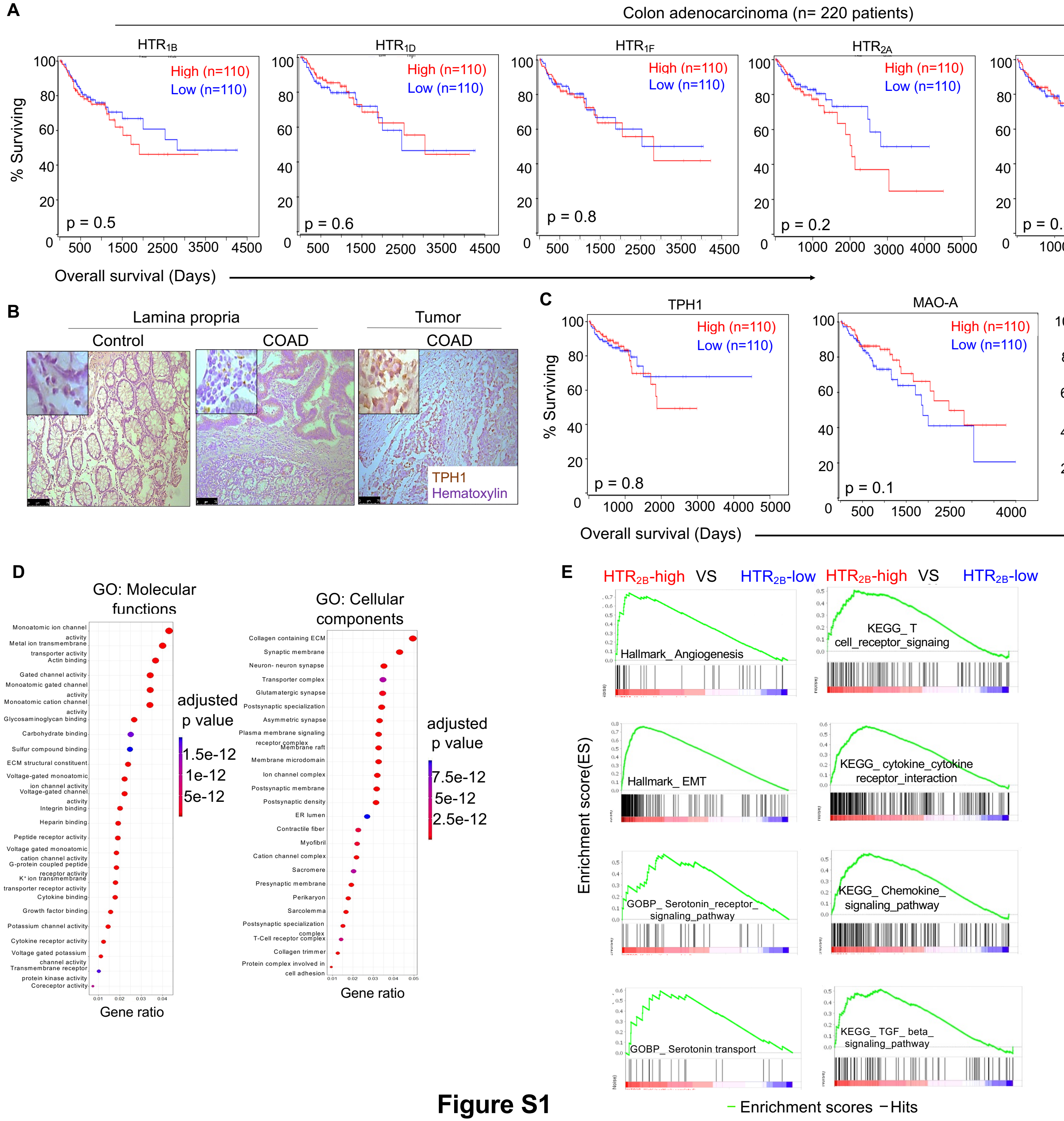


**Figure S1**

**Figure S1: Intra-tumoral $HTR_{2B}$ expression correlates with altered gene expression in colon cancer patients.** **(A)** Kaplan-Meier survival analysis of $HTR_{1B}$, $HTR_{1D}$, $HTR_{1F}$, $HTR_{2A}$, $HTR_{4}$, and $HTR_{7}$ expression(derived from TCGA data) in 220 colon adenocarcinoma patients cohort. The "high" versus "low" gene-expressing data were segregated based on the top or bottom 25% of the cohort. Mantel-cox log-rank test **(B)** Representative images showing the expression of tryptophan hydroxylase 1 (TPH1) in the control (lamina propria) and lamina propria of colon adenocarcinoma (COAD) tissue by immunohistochemistry (IHC) Original magnification 200X, inset images magnification 600X. **(C)** Kaplan-Meier survival analysis of TPH1, MAO-A, and SLC6A expression (derived from TCGA data) in 220 colon adenocarcinoma patients cohort. The "high" versus "low" gene-expressing data were segregated based on the top or bottom 25% of the cohort.,. Mantel-cox log-rank test. p-values were indicated within each plot and p-value less than 0.05 was considered significant. **D-E** RNA-seq datasets from thirty TCGA-COAD samples were segregated into "$HTR_{2B}$ high" and "$HTR_{2B}$ low" (highest fifteen and lowest fifteen intra-tumoral $HTR_{2B}$ expression), and the differential expression of selected genes was analyzed. **(D)** Gene ontology shows altered molecular functions and cellular components from the TCGA-COAD cohort in the "$HTR_{2B}$ high" versus "$HTR_{2B}$ low" datasets (n=15 patients/groups). **(E)** Gene set enrichment analysis (GSEA) of the angiogenesis and epithelial-mesenchymal transition (EMT) hallmark genesets, serotonin receptor signaling pathway and serotonin transport Gene Ontology Biological Processes (GOBP) gene sets, and T cell receptor signaling and cytokine-cytokine receptor interaction and chemokine signaling pathway and TGF beta signaling pathway KEGG pathway genesets. The enrichment score is calculated between the TCGA-COAD cohort from "$HTR_{2B}$ high" versus "$HTR_{2B}$ low" datasets (n=15 patients/groups).

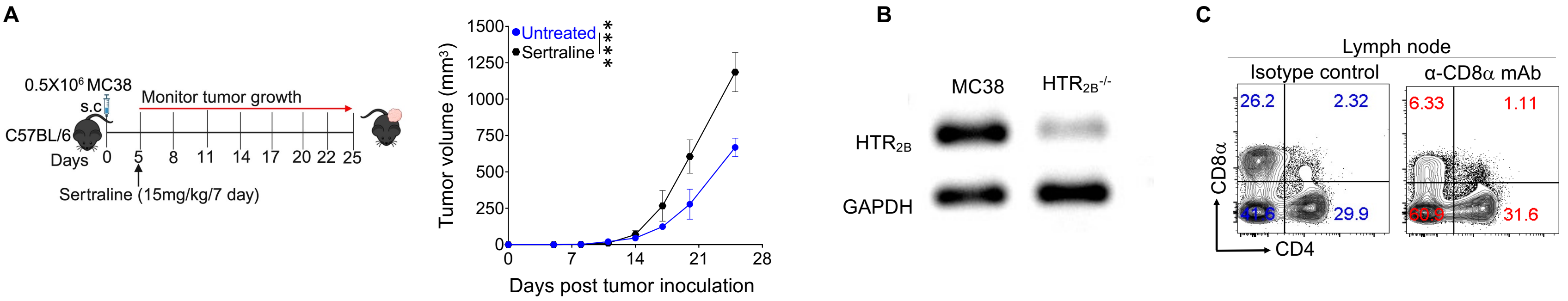


**Figure S2: Higher serum serotonin levels promote tumor progression.** (**A**) C57BL/6 mice were subcutaneously injected with MC38 cells, and from the 5th day of tumor injection, mice were treated with either serotonin reuptake inhibitor (SSRI; sertraline) or left untreated (n=6 mice/group). Tumor volume kinetics are shown. **(B)** Gel images showing the expression of $HTR_{2B}$ mRNA in MC38 cell line before and after transfection with SR-2B CRISPR/Cas9 plasmid. GAPDH is used as an internal control. **(C)** Flow cytometric plot showing the distribution of CD4 and CD8 T cells in tumor-draining lymph nodes from the mice treated with or without CD8 T cell depleting anti-CD8 monoclonal antibody. Two-way ANOVA with Sidak's multiple comparison test (A). ****$p<0.0001$. The error bar represents the ± standard error of means (SEM).

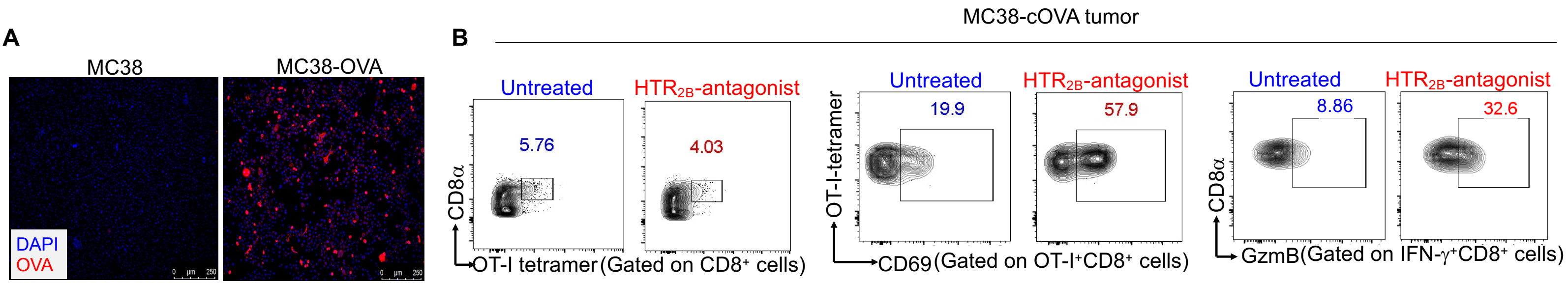


**Figure S3: HTR2B antagonist modulates antigen-specific CD8 T cell response in the tumor**. **(A)** A representative image shows the expression of H-2Kb-SIINFEKL on MC38 cells transfected with pCL-neo-OVA plasmid and MC38 cells without transfection. Original magnification 200X. **(B)** C57BL/6 mice were injected with MC38-cOVA. They were either treated with HTR2B antagonist or left untreated (n=5 mice/ group) for 15 days. On the 15th day, mice were euthanized, and total lymph node cells were tagged with CFSE and restimulated ex vivo with complete ovalbumin protein (50 mg/ml) for 5 days, or immunophenotyping was performed from the cells of the tumor**.** Flow cytometric plots showing the intra-tumoral OT-I tetramer+ total and different subsets of CD8 T cells.
