## Supplementary material for "Antagonizing the serotonin receptor HTR2B drives antigen-specific cytotoxic T-cell responses and controls colorectal cancer growth": Supplemetary Table 1

| **Table 5: Resource table**  Oligonucleotides (Primers) | | | | |
| --- | --- | --- | --- | --- |
| PRIMERS | **SOURCE** | | **GENE IDENTIFIER** | |
| Mouse HTR_1A_ forward  CTGTTCCCTTTGGGTGTCAA | This manuscript | | NM_008308.5 | |
| Mouse HTR_1A_ reverse  AATTCCAGGGCACCATAACC | This manuscript | | NM_008308.5 | |
| Mouse HTR_1B_ forward  TCGAGCAGGGTGATTTCTATTC | This manuscript | | NM_010482.2 | |
| Mouse HTR_1B_ reverse  GTCCTCATTGGACATGGTGTAG | This manuscript | | NM_010482.2 | |
| Mouse HTR_1D_ forward  AGAGTGTCAAAGGCGAGAAC | This manuscript | | NM_001285482.1 | |
| Mouse HTR_1D_ reverse  GCCCAGTGGACACATGATAA | This manuscript | | NM_001285482.1 | |
| Mouse HTR_1F_ forward  GTGAGAGAGAGCTGGATTATGG | This manuscript | | NM_009313.5 | |
| Mouse HTR_1F_ reverse  TAGCCGACAGATGCAAGATG | This manuscript | | NM_009313.5 | |
| Mouse HTR_2A_ forward  TGCTACAACTTCCGGCTTAG | This manuscript | | NM_172812.3 | |
| Mouse HTR_2A_ reverse  CCTCGAGTCGTCACCTAATTG | This manuscript | | NM_172812.3 | |
| Mouse HTR_2B_ forward  TCCTGATACTCGCGGTGATA | This manuscript | | NM_008311.3 | |
| Mouse HTR_2B_ reverse  GTAGCGTACTGCAGCCTTT | This manuscript | | NM_008311.3 | |
| Mouse HTR_2C_ forward  GCCGTCAAACCCTGATGTTA | This manuscript | | NM_008312.4 | |
| Mouse HTR_2C_ reverse  CTCTTCCTCATCACCCTTCTTG | This manuscript | | NM_008312.4 | |
| Mouse HTR_3A_ forward  CAGTATCTTCCTCATGGTCGTG | This manuscript | | NM_013561.2 | |
| Mouse HTR_3A_ reverse  AGACTGAGTATCCCAGAAGGAG | This manuscript | | NM_013561.2 | |
| Mouse HTR_3B_ forward  CTGGAGACTTCGCACAGATT | This manuscript | | NM_020274.4 | |
| Mouse HTR_3B_ reverse  GAGCATCAGGAAGATGCTAGG | This manuscript | | NM_020274.4 | |
| Mouse HTR_4_ forward  CATCCTCTGCTGTGATGATGAG | This manuscript | | NM_008313.4 | |
| Mouse HTR_4_ reverse  TCCCTTAGTACATGGGTGGA | This manuscript | | NM_008313.4 | |
| Mouse HTR_5A_ forward  AGTGAGGAATGCCAAGTCAG | This manuscript | | NM_008314.3 | |
| Mouse HTR_5A_ reverse  CCAGTACACAAAGAGCACCA | This manuscript | | NM_008314.3 | |
| Mouse HTR_5B_ forward  CTACTGGTGTTGCTGATCGT | This manuscript | | NM_010483.3 | |
| Mouse HTR_5B_ reverse  CGAGGCCACCAAGTTATGT | This manuscript | | NM_010483.3 | |
| Mouse HTR_6_ forward  TATTGGCCAGCCTGCCTTA | This manuscript | | NM_001377096.1 | |
| Mouse HTR_6_ reverse  CAAGATCCTGCAGTAGGTGAAG | This manuscript | | NM_001377096.1 | |
| Mouse HTR_7_ forward  CCATCACCTTACCTCCTCTCTT | This manuscript | | NM_008315.3 | |
| Mouse HTR_7_ reverse  GCGGTGGAGTAGATCGTGTA | This manuscript | | NM_008315.3 | |
| Mouse TPH1 forward  GTGGATGACTTTGCGCTAGA | This manuscript | | NM_009414.3 | |
| Mouse TPH1 reverse  CTGAGTCTTGGTGGCTTCTAC | This manuscript | | NM_009414.3 | |
| Mouse DDC forward  CTGGCTGCTCGGACTAAAG | This manuscript | | NM_001190448.1 | |
| Mouse DDC reverse  GCGCCTGATCAGATGTGTAA | This manuscript | | NM_001190448.1 | |
| Mouse Slc6a4 forward  GTGACCAGTGTGGTGAACT | This manuscript | | NM_010484.2 | |
| Mouse Slc6a4 reverse  CACGTCTTCGTTCCTCATCTC | This manuscript | | NM_010484.2 | |
| Mouse MAO-A forward  GAGGTCTTGAATGCTCTAGG | This manuscript | | NM_173740.3 | |
| Mouse MAO-A reverse  TCCTCTCTAAGAAGGTGTGG | This manuscript | | NM_173740.3 | |
| Mouse Cyclophilin-A forward  AGGGTGGTGACTTTACACGC | This manuscript | | NM_011149.2 | |
| Mouse Cyclophilin-A reverse  ATCCAGCCATTCAGTCTTGG | This manuscript | | NM_011149.2 | |
| Mouse GAPDH forward  GGCAAATTCAACGGCACAGT | This manuscript | | NM_017008.4 | |
| Mouse GAPDH reverse  AGATGGTGATGGGCTTCCC | This manuscript | | NM_017008.4 | |
| Antibodies | | | | |
| ANTIBODIES (clone) | **SOURCE** | | **IDENTIFIER** | |
| FITC anti-human CD4 (OKT4) | Biolegend | | Cat# 317408; RRID: AB_571951 | |
| FITC anti-human CD4 (A161A1) | Biolegend | | Cat# 357406; RRID: AB_2562357 | |
| APC anti-mouse CD4 (GK1.5) | Biolegend | | Cat# 100516; RRID: AB_312719 | |
| Biotin anti-mouse CD4 (GK1.5) | Biolegend | | Cat# 100404; RRID: AB_312689 | |
| Biotin anti-mouse/human CD44 (IM7) | Biolegend | | Cat# 103003; RRID: AB_312954 | |
| APC/Cy7 anti-mouse/human CD44 (IM7) | Biolegend | | Cat# 103028; RRID: AB_830785 | |
| PE anti-mouse/human CD44 (IM7) | Biolegend | | Cat# 103008; RRID: AB_312959 | |
| PerCP-Cy5.5 Anti-Mouse CD25 (3C7) | Biolegend | | Cat# 101912; RRID: AB_10613643 | |
| PerCP anti-mouse CD45 (30-F11) | Biolegend | | Cat# 103130; RRID: AB_893339 | |
| PE anti-mouse CD25 (3C7) | Biolegend | | Cat# 101904; RRID: AB_312847 | |
| FITC anti-mouse CD25 (3C7) | Biolegend | | Cat# 101908; RRID: AB_961212 | |
| Biotin anti-mouse CD8a (53-6.7) | Biolegend | | Cat# 100704; RRID: AB_312743 | |
| PerCP-Cy5.5 anti-mouse CD8a (53-6.7) | Biolegend | | Cat# 100734; RRID: AB_2075238 | |
| Alexa Fluor® 700 anti-mouse CD8a (53-6.7) | Biolegend | | Cat# 100730; RRID: AB_493703 | |
| Alexa Flour 647 anti-mouse CD25 (PC61) | Biolegend | | Cat# 102020; RRID: AB_493458 | |
| PE/Cy5 anti-mouse CD25 (PC61) | Biolegend | | Cat# 102010; RRID: AB_312859 | |
| PerCP/Cy5.5 anti-mouse IFN-γ (XMG1.2) | Biolegend | | Cat# 505822; RRID: AB_961359 | |
| PE/Cy7 anti-mouse IFN-γ (XMG1.2) | Biolegend | | Cat# 505826; RRID: AB_2295770 | |
| PE anti-mouse/human T-bet (4B10) | Biolegend | | Cat# 644810; RRID: AB_2200542 | |
| PE/Cy7 anti-mouse/ human T-bet (4B10) | Biolegend | | Cat# 644824; RRID: AB_2561761 | |
| eFlour450 anti-mouse/human T-bet (eBio4B10 (4B10)) | eBioscience | | Cat#48-5825-82; RRID: AB_2784727 | |
| APC anti-mouse CCR7 (4B12) | Biolegend | | Cat# 120108; RRID: AB_389234 | |
| BV711 anti-mouse/human T-bet (4B10) | Biolegend | | Cat# 644820; RRID: AB_2715766 | |
| Spark YG581 Anti-Mouse CD25 (PC61) | Biolegend | | Cat# 102073; RRID: AB_2910275 | |
| PE/Dazzale 594^TM^ anti-mouse PD-L1 (10F.9G2) | Biolegend | | Cat# 124323; RRID: AB_2565638 | |
| BV 421 anti-mouse CTLA-4 (UC10-4B9) | Biolegend | | Cat# 106311; RRID: AB_10901170 | |
| Alexafluor 647 anti-mouse/human granzyme B (GB11) | Biolegend | | Cat# 515405; RRID:AB_2294995 | |
| PE/Cy7 anti-mouse CD62L (MEL-14) | Biolegend | | Cat# 104418; RRID: AB_313103 | |
| Biotin anti-mouse IL-10 (JES5-16E3) | Biolegend | | Cat# 505004; RRID: AB_315358 | |
| PE/Cy7 anti-mouse TNF-α ( N3−19.12) | eBioscience | | Cat# 25-7423-82; RRID: AB_494228 | |
| APC/Cy7 anti-mouse IFN-γ (XMG1.2) | Biolegend | | Cat# 505850; RRID: AB_2616698 | |
| Alexa Fluor 488 anti-serotonin | Novus biologicals | | Cat#NB100-65037AF488; RRID: AB_965390 | |
| Alexa Fluor 647 anti- HTR_2B_ (C6) | Santa-cruz | | Cat#sc-376878; RRID: N/A | |
| Polyclonal purified anti-tryptophan hydroxylase 1 antibody | Novus biologicals | | Cat-NBP1-86922; RRID: AB_11046169 | |
| Alexa Flour 594^™^ anti-mouse CD4 (GK1.5) | Biolegend | | Cat# 100446; RRID: AB_2563182 | |
| Polyclonal goat Anti-Serotonin antibody | Abcam | | Cat#ab66047; RRID: AB_1142794 | |
| Anti-Serotonin transporter antibody (EPR12735) | Abcam | | Cat#ab181034; RRID: AB_2813768 | |
| Alexa Fluor 647^™^ anti-mouse/rat/human FOXP3 (150D) | Biolegend | | Cat# 320014; RRID: AB_439750 | |
| eFlour 506^™^ anti-mouse/human CD44 (IM7) | Thermo fisher scientific | | Cat# 69044182; RRID: AB_2762676 | |
| APC anti-mouse CCR7 (4B12) | Biolegend | | Cat# 120108; RRID: AB_389234 | |
| PE/Cy5.5 anti-mouse FOXP3 (FJK-16s) | Thermo fisher scientific | | Cat# 35577382; RRID: AB_11218094 | |
| APC/Fire 810^™^ anti-mouse PD-1 (29F.1A12) | Biolegend | | Cat# 135251; RRID: AB_2910292 | |
| Biotin anti-mouse CD274 (PD-L1) (10F.9G2) | Biolegend | | Cat# 124305; RRID: AB_961218 | |
| PerCp/Cy5.5 anti-mouse CD274 (PD-L1) (10F.9G2) | Biolegend | | Cat # 124334; RRID: AB_2629832 | |
| Polyclonal Dylight 488 Donkey anti-mouse IgG | Jackson ImmunoResearch | | Cat# 715-486-151; RRID: AB_2572300 | |
| Polyclonal Dylight 488 Donkey anti-rat IgG | Jackson ImmunoResearch | | Cat# 711-486-152; RRID: N/A | |
| Purified anti-mouse CD3ε mAb (145-2C11) | BioXCell | | Cat# BE0001-1, RRID: AB_1107634 | |
| Purified anti-mouse CD28 mAb (37.51) | BioXCell | | Cat# BE0051-1, RRID: AB_1107624 | |
| LEAF™ Purified anti-human CD28 (CD28.2) | Biolegend | | Cat# 302914, RRID: AB_314316 | |
| LEAF™ Purified anti-human CD3 (UCHT1) | Biolegend | | Cat# 300438, RRID: AB_11146991 | |
| Purified IFN gamma mAb (XMG1.2) | eBioscience | | Cat#16-7311-85; RRID: AB_469242 | |
| *InVivo*MAb anti-mouse CD8α (YTS 169.4) | BioXCell | | Cat #BE0117; RRID:AB_10950145 | |
| Rat IgG2b κ isotype control antibody (LTF-2) | BioXCell | | Cat #BE0090; RRID: AB_1107780 | |
| Commercial assay kits | | | | |
| KITS | **SOURCE** | | **IDENTIFIER** | |
| Mouse IFN-γ ELISA MAX^TM^ | Biolegend | | Cat# 430804 | |
| Mouse IL-10 ELISA MAX^TM^ | Biolegend | | Cat# 431414 | |
| TNF-α ELISA MAX Deluxe ELISA kit | Biolegend | | Cat#430904 | |
| Universal Serotonin ELISA Kit | Novus Biologicals | | Cat#NBP2-68127 | |
| Mouse PD-L1 DuoSet ELISA kit | Novus Biologicals | | Cat#DY1019-05 | |
| Cytek® 25-Color Immunoprofiling Assay, cFluor® Reagent Kit (18C) | Cytek Biosciences | | Cat# 900004160 | |
| CellTrace Violet Cell Proliferation Kit | Invitrogen | | Cat#34557 | |
| CellTrace CFSE Cell Proliferation Kit | Invitrogen | | Cat#C1157 | |
| Dynabeads™ Mouse T-Activator CD3/CD28 | Invitrogen | | Cat#11456D | |
| Magnisort^TM^ Mouse CD8 Enrichment Kit | Invitrogen | | Cat#8804-6824-74 | |
| Chemicals, Peptides, plasmids and Recombinant Proteins | | | | |
| REAGENT or RESOURCE | | **SOURCE** | | **IDENTIFIER** |
| Recombinant human IFN gamma | | Biolegend | | Cat# 570202; RRID: N/A |
| Serotonin Hydrochloride | | Santa-Cruz | | Cat# H9523 |
| SB-215505 (HTR_2B_ antagonist) | | Santa Cruz | | Cat# sc-253540 |
| RS-127445 (HTR_2B_ antagonist ) | | Sigma-Aldrich | | Cat# R2533 |
| BW-723C86(HTR_2B_ agonist) | | Sigma-Aldrich | | Cat# B175 |
| Sertraline hydrochloride (Serotonin transporter blocker) | | Abcam | | Cat# ab141068 |
| Pizotifen (pan- HTR_1_, HTR_2A_ and HTR_2C_ antagonist) | | Sigma-Aldrich | | Cat#B9688 |
| SB-269970(HTR_7_ antagonist) | | Sigma-Aldrich | | Cat#S7389 |
| Ketanserin(+) tartarate(pan- HTR_2_ antagonist) | | Sigma-Aldrich | | Cat# S006 |
| Ritanserin(HTR_2A_ antagonist) | | Sigma-Aldrich | | Cat#R103 |
| Cariprazine hydrochloride (HTR_2B_ antagonist) | | Cayman chemicals | | Cat#24025 |
| Phytohemagglutinin (PHA-L) | | eBioscience | | Cat#00-4977-93 |
| Brilliant Violet 421™ H-2K(b) SIINFEKL tetramer | | NIH tetramer core facility | | N/A |
| ViaDye™ Red Fixable Viability Dye Kit | | Cytek Biosciences | | Cat# SKU R7-6008 |
| Propidium iodide | | Biolegend | | Cat# 421301 |
| MTT | | Himedia laboratories | | Cat# TC191 |
| PerCp/Cy5.5 Streptavidin | | Biolegend | | Cat# 405214 |
| Alexafluor 647 Streptavidin | | Biolegend | | Cat# 405237 |
| PE/Cy7 Streptavidin | | Biolegend | | Cat# 405206 |
| Dylight 549 Streptavidin | | Jackson ImmunoResearch | | Cat# 016-500-084 |
| Dylight 649 Streptavidin | | Jackson ImmunoResearch | | Cat# 016-490-084 |
| pCl-neo-cOVA | | Addgene | | Plasmid #25097 |
| Lipofectamine 3000 | | Thermo Fisher Scientific | | Cat#L3000001 |
| SR-2B CRISPR/Cas9 KO plasmid | | Santa Cruz biotechnology | | Cat# sc-420994 |
| Retrieve-All Antigen Unmasking: Universal 1X | | Biolegend | | Cat#928001 |
| 3,3'- Diaminobenzidine tetrahydrochloride hydrate (DAB) | | Sigma-Aldrich | | Cat# D5637 |
| H_2_O_2_ | | Fisher Scientific | |  |
| Lymphoprep | | Stem Cell technologies | | Cat# 07811 |
| Media and Buffers | | | | |
| Cell Stimulation cocktail | | 50ng/ml Phorbol 12-myristate 13-mcetate (PMA) (MP Biomedicals; Cat# 0215186401)  750ng/ml Ionomycin (MP Biomedicals; Cat# 0215961101)  1X Brefeldin A (Biolegends; Cat# 420601) | |  |
| Tissue digestion reagents | | 0.2mg/ml Dnase I (MP Biomedicals; Cat# 0210057583)  0.1mg/ml Collagenase IV (MP Biomedicals; Cat# 195110)  0.1mg/ml Collagenase I (MP Biomedicals; Cat# 195109)  0.1mg/ml Collagenase D (Sigma; Cat# 11088858001)  0.25mg/ml Dispase I (Roche; Cat# 04942086001)  0.1mg/mg Trypsin inhibitor (Sigma; Cat# T6522)  0.06mg/ml Hyaluronidase (Sigma; Cat# H3506) | |  |
| 1X PBS | | 8 gram/litre NaCl (Fisher Scientific)  0.2 grams/litre of KCl (Sigma-Aldrich)  1.44 gram/litre of Na_2_HPO_4_ (Sigma-Aldrich)  0.24 gram/litre of KH_2_PO_4_ (Fisher Scientific) | |  |
| Flow cytometry cell staining buffer | | 1X PBS  2%BSA(w/v) (Himedia; Cat# TCL146) | |  |
| Complete DMEM (Growth media for MC38 cell culture) | | 13.4 gram/litre DMEM with 4.5 gram/litre D-Glucose and 4 mM L-Glutamine (Himedia; Cat# AT066)  3 gram/litre NaHCO_3_ (Fisher Scientific)  10% FBS (GIBCO/ MP Biomedicals)  1 mM Sodium Pyruvate (Invitrogen)  1 mM non-essential amino acid (Invitrogen)  0.05mg/ml gentamycin (Invitrogen)  100 U/ml Penicillin (Invitrogen)  100 µg/ml Streptomycin (Invitrogen)  pH was adjusted to 7.4 | |  |
| Complete RPMI (Growth media for CD4 T cell culture) | | 16.4 gram/litre RPMI with 4.5 gram/litre D-Glucose and 4 mM L-Glutamine  (Himedia Laboratories/Invitrogen)  1.5 gram/litre NaHCO_3_ (Fisher Scientific)  10% FBS (GIBCO/ MP Biomedicals)  1 mM Sodium Pyruvate (Invitrogen)  100 U/ml Penicillin (Invitrogen)  100 µg/ml Streptomycin (Invitrogen)  pH was adjusted to 7.4 | |  |
| Complete RPMI (Growth media for 4T1-Luc2 cell culture) | | 16.4 gram/litre RPMI with 4.5 gram/litre D-Glucose and 4 mM L-Glutamine  (Himedia; Cat# AT060)  1.5 gram/litre NaHCO_3_ (Fisher Scientific)  10% FBS (GIBCO/ MP Biomedicals)  1 mM Sodium Pyruvate (Invitrogen)  100 U/ml Penicillin (Invitrogen)  100 µg/ml Streptomycin (Invitrogen)  pH was adjusted to 7.4 | |  |
| Cell Freezing medium (For MC38, 4T1-Luc2 and B16F10) | | Complete RPMI 1640 (for 4T1-Luc2) / DMEM (for MC28 and B16F10)  25 % FBS (GIBCO/ MP Biomedicals)  5% DMSO (MP Biomedicals) | |  |
| ACK lysis buffer | | 8.26 gram/litre NH_4_Cl (Fisher Scientific)  1 gram/litre KHCO_3_ (Himedia laboratories)  37.2 milligram/litre Na_2_.EDTA (Fisher Scientific) | |  |
| Cell lines | | | | |
| Cell Lines | | **Source** | | **Identifier** |
| MC38 | | Kerafast | | ENH204-FP |
| 4T1-Luc2 | | ATCC | | CRL-2539-LUC2 |
| MC38-cOva | | This manuscript | |  |
| B16F10 | | NCCS repository | | CRL-6475 |
| Experimental Models: Organisms/ Strains | | | | |
| Mice strains | | **SOURCE** | | **IDENTIFIER** |
| C57BL/6J | | The Jackson Laboratory (Bar Harbor, ME) | | RRID: IMSR_JAX:000664 |
| BALB/CJ | | The Jackson Laboratory | | RRID: IMSR_JAX:000651 |
| C57BL/6-Tg (TcraTcrb)1100Mjb/J  (OT-I) | | The Jackson Laboratory | | RRID: IMSR_JAX:003831 |
| B6.SJ: L-Ptprc^a^ Pepc^b^/BoyJ  (B6.CD45.1) | | The Jackson Laboratory | | RRID: IMSR_JAX:002014 |
| NOD.Cg-*Rag1^tm1Mom^* *Il2rg^tm1Wjl^*/SzJ  (NRG mice) | | The Jackson Laboratory | | RRID: IMSR_JAX:007799 |
| Software and Algorithms | | | | |
| SOFTWAREs | | **SOURCE** | | **IDENTIFIER** |
| Leica MMAF software | | Leica | | <https://www.leica-microsystems.com/> |
| CellSens software | | Olympus | | https://www.olympus-lifescience.com/en/software/cellsens/ |
| Image-J software | | NIH, USA | | <https://imagej.nih.gov/ij/> |
| GraphPad Prism 10 | | GraphPad | | [https://www.graphpad.com](https://www.graphpad.com/) |
| FlowJo v10.2 | | Tree Star | | [https://www.ﬂowjo.com/](https://www.flowjo.com/) |
| Spectroflow | | Cytek Biosciences | | https://spectrum.cytekbio.com/ |
| OMIQ | | Dotmatrics | | https://www.omiq.ai/ |
| FCS Express 7 | | DeNovo Lifesciences | |  |
| R studio | | Posit | |  |
| GSEA | | Broad Institute | |  |
| Morpheus | | Broad Institute | | https://software.broadinstitute.org/morpheus |
| Endnote | | Clarivate | | https://endnote.com/ |
